# Migration genetics link excitatory neurons to ancient selection and economic growth

**DOI:** 10.64898/2026.02.05.703995

**Authors:** Lucas G. Casten, Asthon Tener, Muhammad Elsadany, Ji Seung Yang, John F. Strang, Jacob J. Michaelson

## Abstract

From ancient nomadic movements to modern urbanization, migration has driven human history through mechanisms that remain poorly understood. We conducted a genome-wide association study of migration distance in 250,000 UK individuals, identifying 20 loci in neurodevelopmental genes with 5% heritability. Migration-related variants are associated with excitatory neuron gene expression and correlate with cognition, risk-tolerance, and reduced interpersonal attachment. Within-family analyses demonstrate genetic effects remain significant after controlling for shared environmental factors between siblings. Our polygenic score predicts inferred mobility in *>*1,000 ancient individuals spanning 10,000 years, revealing positive selection on migration alleles that increased substantially over millennia. At the population level, each standard deviation increase in county-level migration polygenic score predicts *>*$4,000 greater income growth per person in the US. These findings establish migration as a heritable trait and suggest biological pathways connecting individual neurodevelopment with regional prosperity across evolutionary and contemporary timescales.

---

Migration has “always been… one of the primary sources of progress”, placing people in sustained contact with those unlike themselves (*1*). From the initial expansion out of Africa to modern urbanization, the movement of people has reshaped languages, economies, and gene pools at every scale (*2–4*). Migrants have consistently advanced frontiers by adapting and innovating across cultural barriers, and their communities reap measurable economic benefits (*5*). Recent years have seen unprecedented global population movement driven by economic opportunity, conflict, climate change, and politics (*6, 7*). Despite intensifying public debate about who migrates and why, the science of individual migration propensity remains in its infancy; we lack a fundamental understanding of why individuals differ in their propensity to move.

Prevailing theoretical models of migration focus heavily on predisposing “push-pull” circumstances, casting individuals as actors responding uniformly and rationally to external conditions (*8, 9*). However, these models struggle to explain substantial variation in migration behavior among individuals facing identical circumstances, including within families where one sibling migrates while another remains. This gap points to poorly characterized “personal factors” that may include innate, biological influences on the urge to migrate. Given migration’s central role in shaping human evolution and contemporary societies, understanding individual differences in migration propensity could inform questions spanning evolutionary biology, neuroscience, and social policy.

While genetics has proven powerful for reconstructing humanity’s ancient migrations (*2, 3, 10*), relatively few studies have investigated whether genetic variation causally influences individual migration propensity in contemporary populations. A recent genome-wide association study identified genetic variants associated with migration distance in the UK, demonstrating significant SNP-based heritability and positive genetic correlations with educational attainment and health outcomes (*11*). These results support theories that migration selects for individuals with greater skills and better health. Polygenic score approaches have validated the links between the genetic architecture for education and migration (*12, 13*). However, recent methodological advances have revealed that GWAS of behavioral traits can be substantially confounded by social stratification: individuals sort into different geographic regions and socioeconomic strata based partly on heritable traits, creating spurious associations between genetic variants and environmental exposures (*14, 15*). Critical questions thus remain: Do migration genetics reflect direct biological influences or are they simply consequences of this socioeconomic sorting? What biological mechanisms and cell types underlie these associations? How consistent are genetic influences across families, ancestries, and evolutionary time? Does the heritable component of migration predict migration’s documented economic benefits at the population level?

Here we integrate multiple lines of evidence to comprehensively characterize the genetic architecture of human migration, its biological mechanisms, evolutionary origins, and societal consequences. We conduct genome-wide association analysis in approximately 250,000 UK Biobank participants (*16*), establishing both between-family and within-family genetic effects to separate genetic influences from those arising through gene-environment correlations. Functional genomic analyses use spatial transcriptomics, single-cell RNA sequencing, and cell-type-specific eQTL mapping to identify the cellular mechanisms through which migration-associated variants act. Validation across an independent US cohort and general population samples spanning five ancestral superpopulations and sibling pairs establishes the robustness of genetic influences across diverse populations and family structures. Ancient DNA analysis spanning 10,000 years tests whether migration genetics predict mobility across human evolutionary history and reveals signatures of positive selection. Finally, county-level analysis across the United States (using a sample enriched for neurodiversity and LGBTQ+) examines whether population-level shifts in migration genetics predict regional economic growth, independent of educational effects.

Our findings reveal that migration propensity manifests through early neurodevelopmental processes in the prenatal brain, influencing gene expression in cortical excitatory neurons rather than reflecting population stratification artifacts. Migration associates with a behavioral profile marked by increased cognitive performance, risk-tolerance, openness, and reduced interpersonal attachment. The same genetic variants that predict modern migration also predict mobility in ancient individuals spanning 10,000 years, from Bronze Age nomadic movements to modern intercontinental relocations, and show signatures of positive natural selection. At the population level, county-level shifts in migration-associated genetic variation predict regional economic growth beyond the effects of educational attainment alone. These results establish migration as an evolutionarily consequential trait whose effects span individual neurodevelopment to regional economic outcomes.

## Results

### Migration is associated with education, health, and independence across cultures

We operationalized migration as the geographic distance between an individual’s birthplace and their current (or final) location, assembling migration distance measures across three populations: approximately 390,000 European-ancestry UK Biobank participants (*16*), *>* 3,000 US adults from the SPARK cohort (*17*) (ascertained for autism research; 70.2% with ASD and 52.3% identifying as LGBTQ+), and approximately 1,300 ancient individuals spanning roughly 10,000 years (*18, 19*). Migration distance correlated with education, health, and independence in both the UK discovery (max N = 393,544) and US replication (max N = 3,235) samples. Mean migration distances differed between cohorts (UK: mean = 47 miles, median = 11 miles; US: mean = 345 miles, median = 33 miles), yet phenotypic correlation patterns were consistent across populations.

Educational attainment showed similar positive correlations with migration distance in both the UK (*r* = 0.26, *p <* 2 × 10^−16^, N = 390,390; Figure 2C) and US (*r* = 0.2, *p <* 2 × 10^−16^, N = 3,010; Figure 2G). Phenotypic analysis revealed that migration distance in the UK positively correlated with cognitive performance (fluid intelligence: *β* = 0.21, FDR *<* 2 × 10^−16^), career mobility (number of jobs held: *β* = 0.051, FDR = 7.9 × 10^−14^), sexual behavior (lifetime sexual partners: *β* = 0.031, FDR *<* 2 × 10^−16^), fertility outcomes in women (age at first live birth: *β* = 0.19, FDR *<* 2 × 10^−16^; number of live births: *β* = -0.026, FDR *<* 2 × 10^−16^), and feelings about work outcomes (work satisfaction: *β* = 0.025, FDR = 8.7 × 10^−12^; financial satisfaction: *β* = 0.06, FDR *<* 2 × 10^−16^). Conversely, migration distance negatively correlated with neuroticism (*β* = -0.046, FDR *<* 2 × 10^−16^), interpersonal relationships (friend satisfaction *β* = -0.06, FDR *<* 2 × 10^−16^; family satisfaction *β* = -0.037, FDR *<* 2 × 10^−16^), and socioeconomic deprivation (Townsend index: *β* = -0.096, FDR *<* 2 × 10^−16^). Health-related measures showed migration distance positively associated with subjective health satisfaction (*β* = 0.029, FDR = 1 × 10^−15^) and physical fitness markers (hand grip strength: *β* = 0.027, FDR *<* 2 × 10^−16^; basal metabolic rate: *β* = 0.01, FDR *<* 2 × 10^−16^). Migration distance negatively correlated with average blood sugar levels (A1C: *β* = -0.03, FDR *<* 2 × 10^−16^), inflammatory markers (C-reactive protein: *β* = -0.037, FDR *<* 2 × 10^−16^), medication use (*β* = -0.05, FDR *<* 2 × 10^−16^), and adiposity measures (body fat percentage: *β* = -0.073, FDR *<* 2 × 10^−16^; BMI: *β* = -0.08, FDR *<* 2 × 10^−16^).

**Figure 1:**
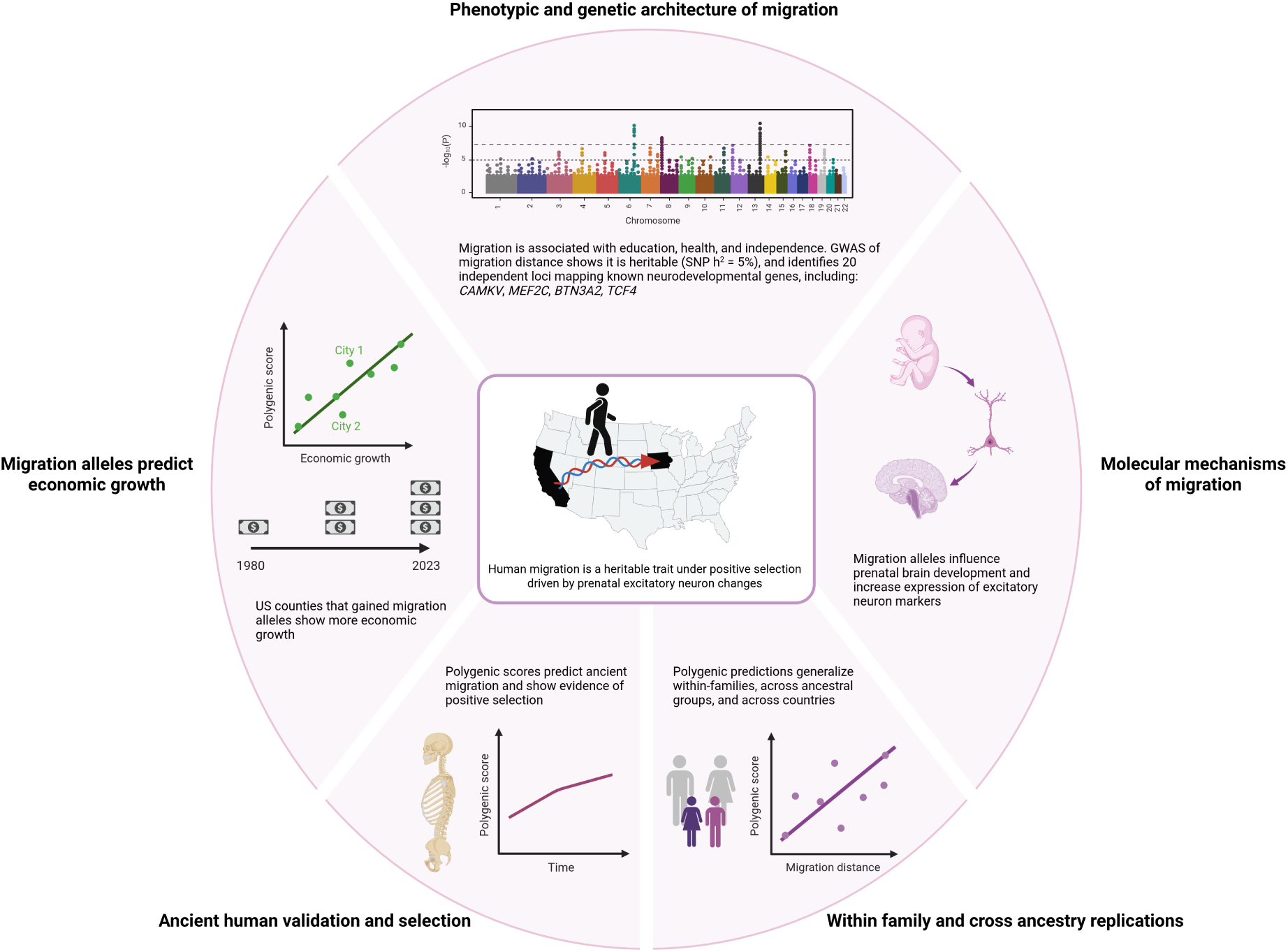
Overview of this study and key results.

**Figure 2:**
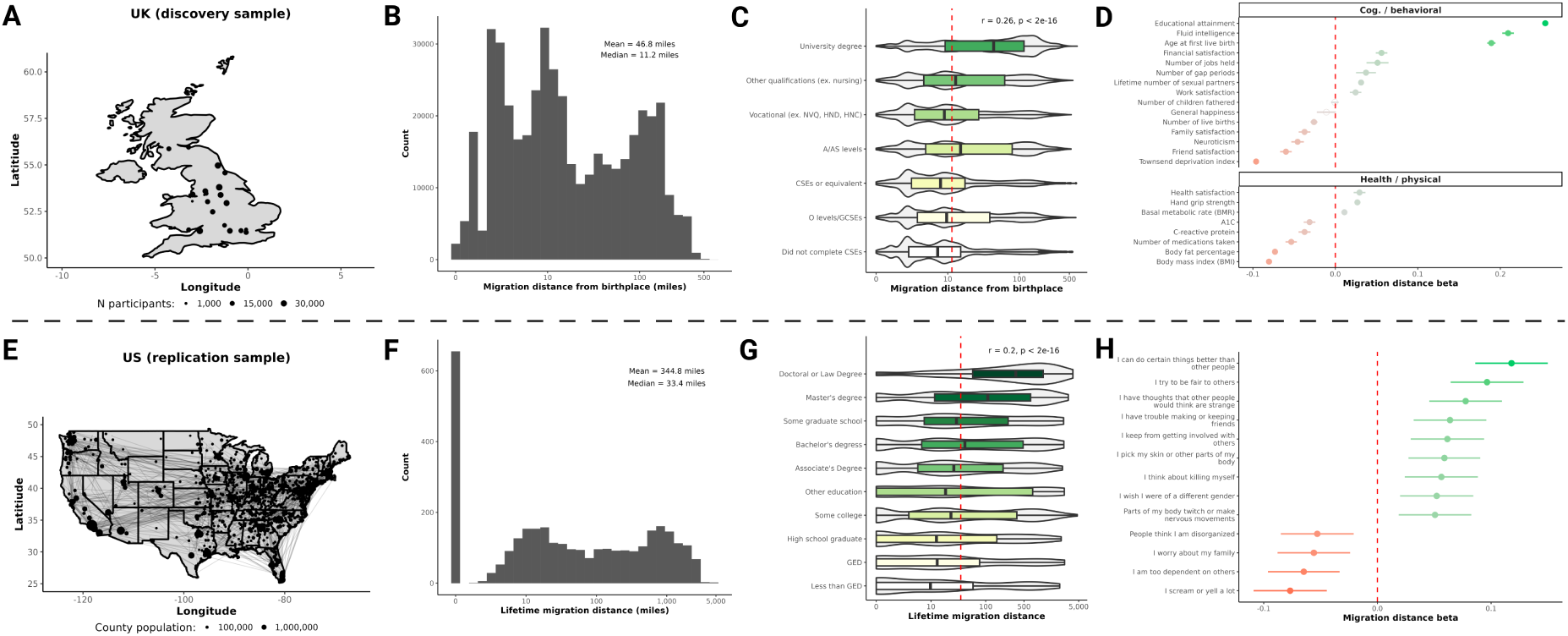
Migration is associated with education, health, and interpersonal relationships in the UK and US. (**A**) Map showing the locations UK Biobank sites and the number of discovery samples used in the study. (**B**) Histogram showing the distribution in miles of migration distances in the UK Biobank. Mean and median migration distances are also reported. (**C**) Distribution of migration distances across educational qualifications in the UK Biobank are shown with violin and boxplots. Pearson correlation statistics between migration distance and years of education (based on these qualifications) are shown. (**D**) Forest plot showing the association between migration distance and phenotypes related to cognition, socioeconomic status, personality, and health are shown from the UK Biobank. Points represent linear regression *β* estimates and 95% confidence intervals. Green colors indicate strong positive associations between migration and the trait, while red indicate negative. Sample sizes varied for each trait, full summary statistics are reported in Supplemental Table 1. (**E**) Map showing the counties where US sample participants live, lines connecting counties indicate where these participants moved to and from. Size of the points are indicative of the 2023 population size for each county. (**F**) Histogram showing the distribution in miles of migration distances in the US sample. Mean and median migration distances are also reported. (**G**) Distribution of migration distances across educational qualifications in the US sample are shown with violin and boxplots. Pearson correlation statistics between migration distance and years of education (based on these qualifications) are shown. (**H**) Forest plot showing the association between migration distance and phenotypes related to self-reported traits captured with the Adult Self Report questionnaire (N = 3,117) in the US sample. Points represent linear regression *β* estimates and 95% confidence intervals. Green colors indicate strong positive associations between migration and the trait, while red indicate negative. Only items with a FDR *<* 0.05 are shown. Full summary statistics are reported in Supplemental Table 2.

The US sample demonstrated convergent patterns, with migration distance positively correlating with self-reported capabilities (“I can do certain things better than other people”, *β* = 0.12, FDR = 6.4 × 10^−11^), and negatively correlating with dependence on others (“I am too dependent on others”, *β* = -0.065, FDR = 0.001) as well as friend relationships (“I have trouble making or keeping friends”: *β* = 0.064, FDR = 0.002) and familial relationships (“I get along badly with my family”, *β* = 0.063, FDR = 0.003). These findings from an autism-ascertained sample strengthen confidence in the robustness of migration-personality associations across diverse populations. Together, the cross-cultural consistencies indicate migratory behavior captures a broader phenotypic profile characterized by higher general cognitive abilities, greater independence, and better physical health, alongside less satisfaction with social relationships.

### GWAS of migration reveals genetic correlations with cognition, personality, and health

To elucidate the genetic architecture underlying migratory behavior, we conducted a genomewide association study (GWAS) of migration distance in unrelated European ancestry individuals from the UK Biobank (N = 242,561). We identified 20 genome-wide significant independent loci (*p <* 5 × 10^−8^; Supplemental Table 3; Figure 3A), with lead variants mapping to genes previously implicated in neurodevelopment and neuropsychiatric phenotypes, including *NEGR1*, *CAMKV*, *BCL11A*, *MEF2C*, *BTN3A2*, *PCDH17*, and *TCF4* (*20–25*). SNP-based heritability (*h*^2^) was estimated at 4.97% (SE = 0.37%, *p <* 2 × 10^−16^), indicating that common genetic variants contribute significantly but modestly to individual differences in migration distance.

**Figure 3:**
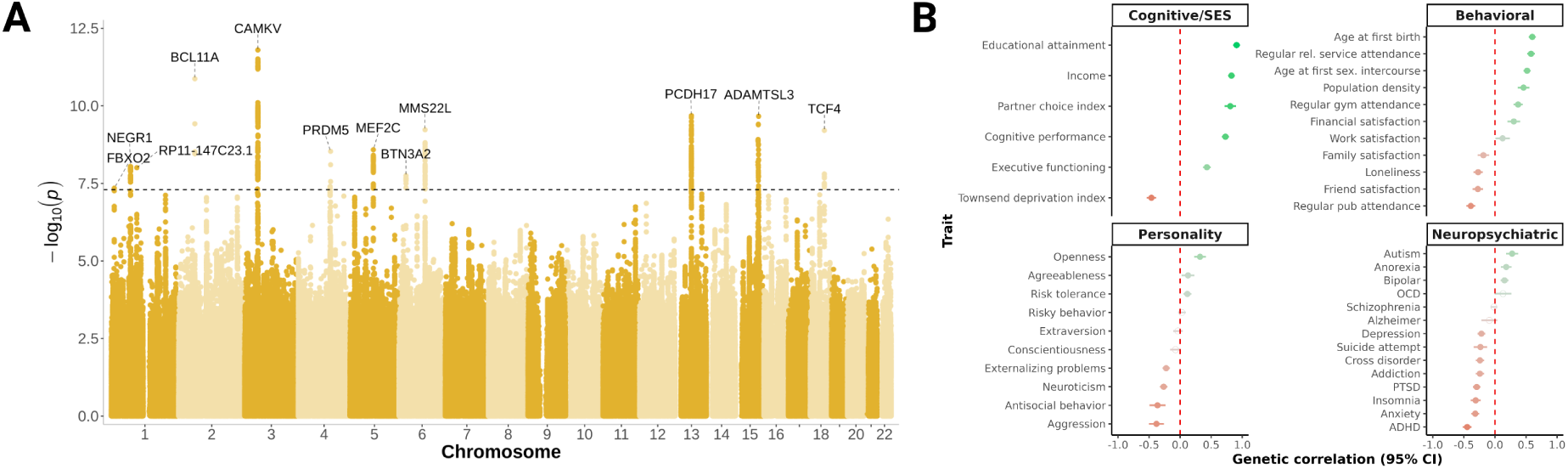
Genetic associations of migration distance. (**A**) Manhattan plot of GWAS summary statistics for migration distance in the UK Biobank. Lead SNPs with a *p <* 5×10^−8^ are annotated with the nearest gene based on hg19 coordinates. (**B**) Genetic correlations (**r*_*g*_*) estimated using LDSC between migration distance and various brain and behavioral traits are shown. Points represent **r*_*g*_* estimates and 95% confidence intervals. Green colors indicate strong positive correlations between migration and the trait, while red indicate negative. Full summary statistics can be found in Supplemental Table 5.

To assess shared genetic architecture between migration distance and other traits, we calculated genetic correlations (*r_*g*_*) using Linkage Disequilibrium Score Regression (LDSC; 3B) (*26*). Migration distance showed significant positive genetic correlations with cognitive and socioeconomic traits (educational attainment: *r_*g*_* = 0.91, FDR = 5.2 × 10^−217^; income: *r_*g*_* = 0.83, FDR = 6.7 × 10^−196^; cognitive performance: *r_*g*_* = 0.73, FDR =2.2 × 10^−140^) and reproductive measures (age at first birth: *r_*g*_* = 0.6, FDR = 9.3 × 10^−98^; age at first sexual intercourse: *r_*g*_* = 0.52, FDR = 2.3 × 10^−71^). Personality traits associated with migration included openness (*r_*g*_* = 0.32, FDR = 6.4 × 10^−11^) and risk tolerance (*r_*g*_* = 0.12, FDR = 3.9 × 10^−4^), while showing negative genetic correlations with aggression (*r_*g*_* = -0.38, FDR = 8.1 × 10^−10^), neuroticism (*r_*g*_* = -0.26, FDR = 1.3 × 10^−16^), and externalizing problems (*r_*g*_* = -0.22, FDR = 7.2 × 10^−13^).

Genetic correlations with neuropsychiatric conditions were heterogeneous in direction. Migration distance showed positive genetic correlations with autism (*r_*g*_* = 0.28, FDR = 4.8 × 10^−8^) and bipolar disorder (*r_*g*_* = 0.15, FDR = 3.5×10^−6^), but negative correlations with many other diagnoses, including: ADHD (*r_*g*_* = -0.44, FDR = 1.4 × 10^−33^), anxiety (*r_*g*_*= -0.32, FDR = 2.1 × 10^−22^), and depression (*r_*g*_* = -0.22, FDR = 3.2 × 10^−12^).

Given the high genetic correlation with educational attainment, we performed GWAS-by-subtraction to verify that migration captures genetic variation independent of education (*27*). The education-independent component remained significantly heritable (*h*^2^ = 1.9%, Z = 3.2) and identified multiple independent genome-wide significant loci (including one in *HOMER2*), confirming that migration reflects distinct genetic influences beyond education (Figure S1; Supplemental table 7). The education-independent component showed positive genetic correlations with cognitive performance and risk-taking, paralleling the phenotypic associations reported above (Supplemental table 8). The extensive genetic overlap reveals that pleiotropic genetic variants simultaneously influence migration, cognition, personality, and mental health through shared neurodevelopmental mechanisms.

### Migration-associated variants are associated with gene expression in cortical excitatory neurons

To elucidate the molecular and cellular mechanisms underlying heritable variation in human migration, we conducted functional genomic analyses of our UK Biobank migration GWAS results through three complementary approaches: spatial transcriptomics, single-cell RNA sequencing, and cell-type-specific expression quantitative trait loci (eQTL) mapping.

We first sought to localize the tissue(s) where migration-related genetic variation is enriched. We utilized a recently developed method (gsMap (*28*)) that integrates GWAS summary statistics with spatial transcriptomics to identify tissues and regions with enriched heritability. To examine an unbiased set of tissues, we applied this approach to whole mouse embryo transcriptomic data (*29*). Migration heritability was most enriched in brain tissue (*p* = 8.3 × 10^−8^; Figure 4D). Minimal enrichment was observed in non-brain tissues, indicating that migration-associated variants primarily influence neurobiology.

**Figure 4:**
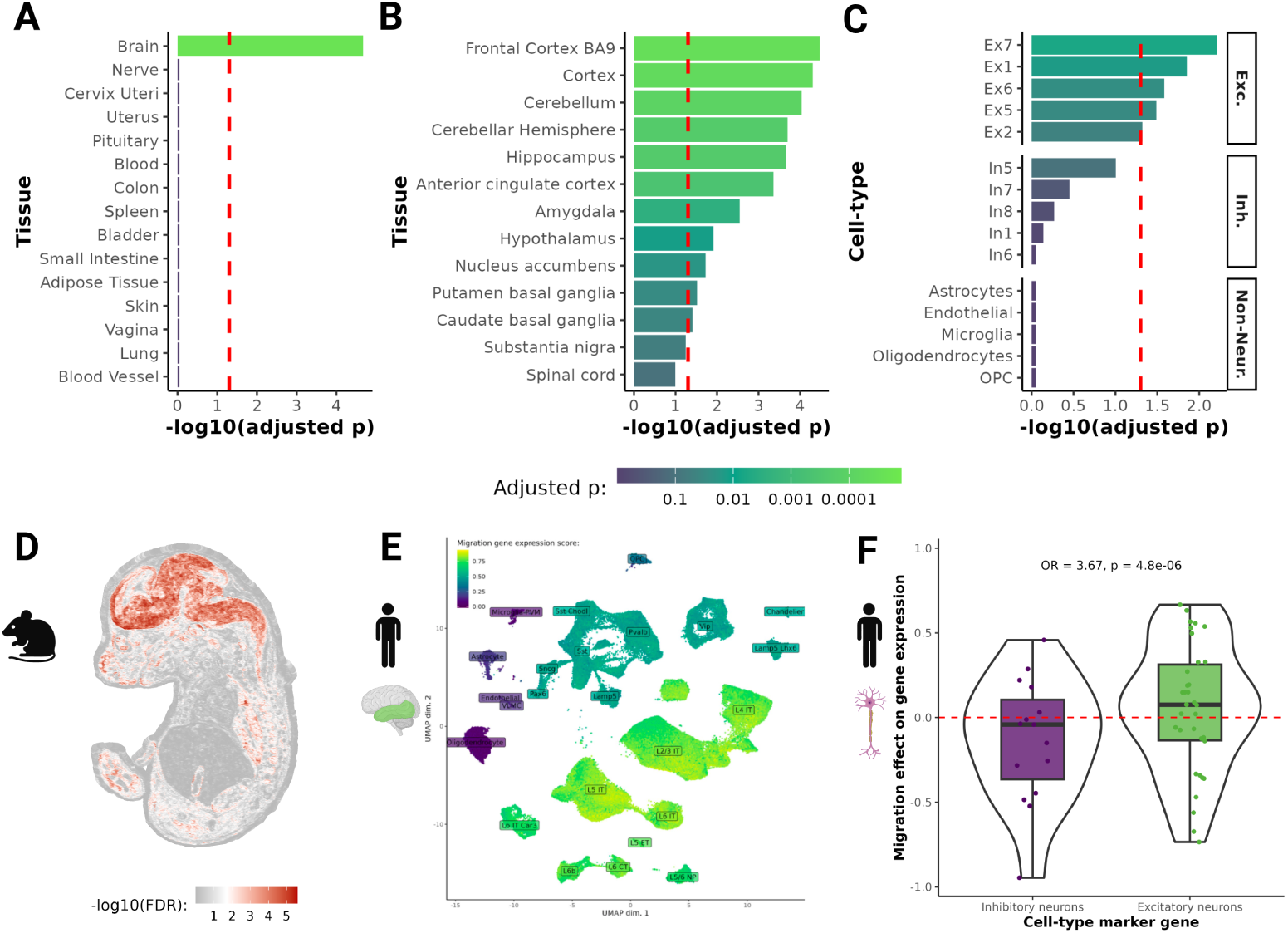
Migration variants influence excitatory neuron function. (**A**) Enrichment for migration-related variation across GTEx v8 tissues computed using MAGMA, top 15 tissues are shown. Bonferroni adjusted *p* across all 30 tissues is shown. (**B**) Enrichment for migration-related variation across all brain GTEx v8 tissue subtypes computed using MAGMA. Bonferroni adjusted *p* across all 53 GTEx tissue subtypes is shown. (**C**) Enrichment for migration-related variation across PsychENCODE development cell-types computed using MAGMA. Top 5 sub-types for excitatory neurons (“Exc.”), inhibitory neurons (“Inh.”), and non-neuronal (“Non-Neur.”) are shown. Bonferroni adjusted *p* across all cell-types is shown. (**D**) Spatial mapping of migration-associated variants across a developing mouse embryo (E16.5 days). Results are based on the gsMap method, which tests for heritability enrichment across spatial transcriptomic data. Red indicates enrichment of migration related genes. (**E**) Cell-type specific mapping from the scPagwas method, mapping migration-associated variants across postmortem human brain tissue from the medial temporal cortex. Green indicates higher expression of migration associated genes. X-axis represents UMAP dimension 1 and Y-axis represents UMAP dimension 2. Cell-type labels are overlaid on the center of each cluster. (**F**) Distribution of migration-gene expression correlations for marker genes of excitatory (green) and inhibitory (purple) neurons. Each point represents a cell-type marker gene. Values represent the correlation between migration GWAS and single-cell eQTL effect sizes in excitatory neurons. Positive values indicate tha_2_t _3_migration-increasing alleles are associated with increased gene expression.

To identify the specific cell types mediating these neurodevelopmental effects, we performed cell-type-specific enrichment analysis using single-cell RNA sequencing data from human postmortem brain tissue. We applied the scPagwas method, which links GWAS variants to cell-typespecific gene expression programs (*30*). Migration distance showed significant heritability enrichment across all excitatory neuron subtypes (Figure 4E), with the strongest enrichment observed in cortical excitatory populations layer 5 intratelencephalic (*p* = 1.1 × 10^−8^), layer 6 intratelencephalic (*p* = 2.8 × 10^−8^, and layer 2/3 intratelencephalic neurons (*p* = 2.8 × 10^−7^)). In contrast, we observed no significant enrichment in inhibitory neurons subtypes or non-neuronal cells.

To determine the direction of these effects, we correlated SNP effects on migration distance with their effects on gene expression using single-cell eQTL data from both excitatory and inhibitory neurons in human brain tissue. We focused our analysis on a set of cell-type marker genes from the DESCARTES atlas (*31*), selecting high-confidence markers based on expression specificity (TPM *>* 100, fold change *>* 1.5, and adjusted *p <* 0.01) for conservative cell-type marker gene assignment. For each marker gene, we calculated the correlation between migration GWAS effect sizes and scQTL effect sizes (*32*), quantifying the direction of gene expression changes associated with migration-increasing alleles. Migration-associated variants were significantly more likely to show positive correlations with marker gene expression in excitatory neurons compared to inhibitory neurons (odds ratio = 3.67, *p* = 4.8 × 10^−6^; Figure 4F), indicating that migration-increasing alleles disproportionately upregulate excitatory neuron marker genes. We confirmed this cell-type-specific pattern using a complementary approach that directly compared the correlation between migration effects and scQTL effects within individual genes across both cell types (N = 116 neuronal markers from PanglaoDB (*33*)). This analysis revealed that 23% of genes exhibited excitatory neuron-specific effects (migration alleles increased expression in excitatory neurons but decreased expression in inhibitory neurons) versus only 9% with inhibitory neuron-specific effects (excitatory/inhibitory enrichment = 2.9, *p* = 0.008; Figure 4G). These cell-type-specific patterns indicate that migration-associated variation disproportionately increases excitatory neuron marker gene expression relative to inhibitory neuron expression.

We validated the observed enrichment in brain tissues and excitatory neurons using MAGMA-based (*34, 35*) gene-set analysis in independent datasets, yielding concordant results (Figure 4A-C, Supplementary Figure S5). Collectively, these findings indicate that genetic variation influencing human migration is associated with gene expression in cortical excitatory neurons, a group of neurons involved in higher-order cognitive processing and long-range cortical projections.

### Within family and cross-population validation of the migration polygenic score

To evaluate the cross-population validity of our genetic findings and to establish causality independent of population stratification and indirect genetic effects, we tested whether a polygenic score (PGS) derived from our GWAS in individuals of White British European ancestry could predict migration behavior in independent samples, populations, and within sibling pairs. These analyses allow us to determine whether genetic effects on migration persist across diverse populations and familial environments.

The migration PGS significantly predicted migration distance across all five ancestral superpopulations in the UK Biobank, including European (EUR; N = 22,301), African (AFR; N = 2,237), Admixed American (AMR; N = 1,232), East Asian (EAS; N = 961), and South Asian (SAS; N = 793) ancestry groups (Figure 5A). Effect sizes were highly consistent across populations (range: β = 0.11 to 0.18), suggesting genetic portability despite the PGS being derived from European ancestry individuals. This cross-population consistency suggests that the genetic architecture underlying migration propensity is shared across diverse ancestral backgrounds.

**Figure 5:**
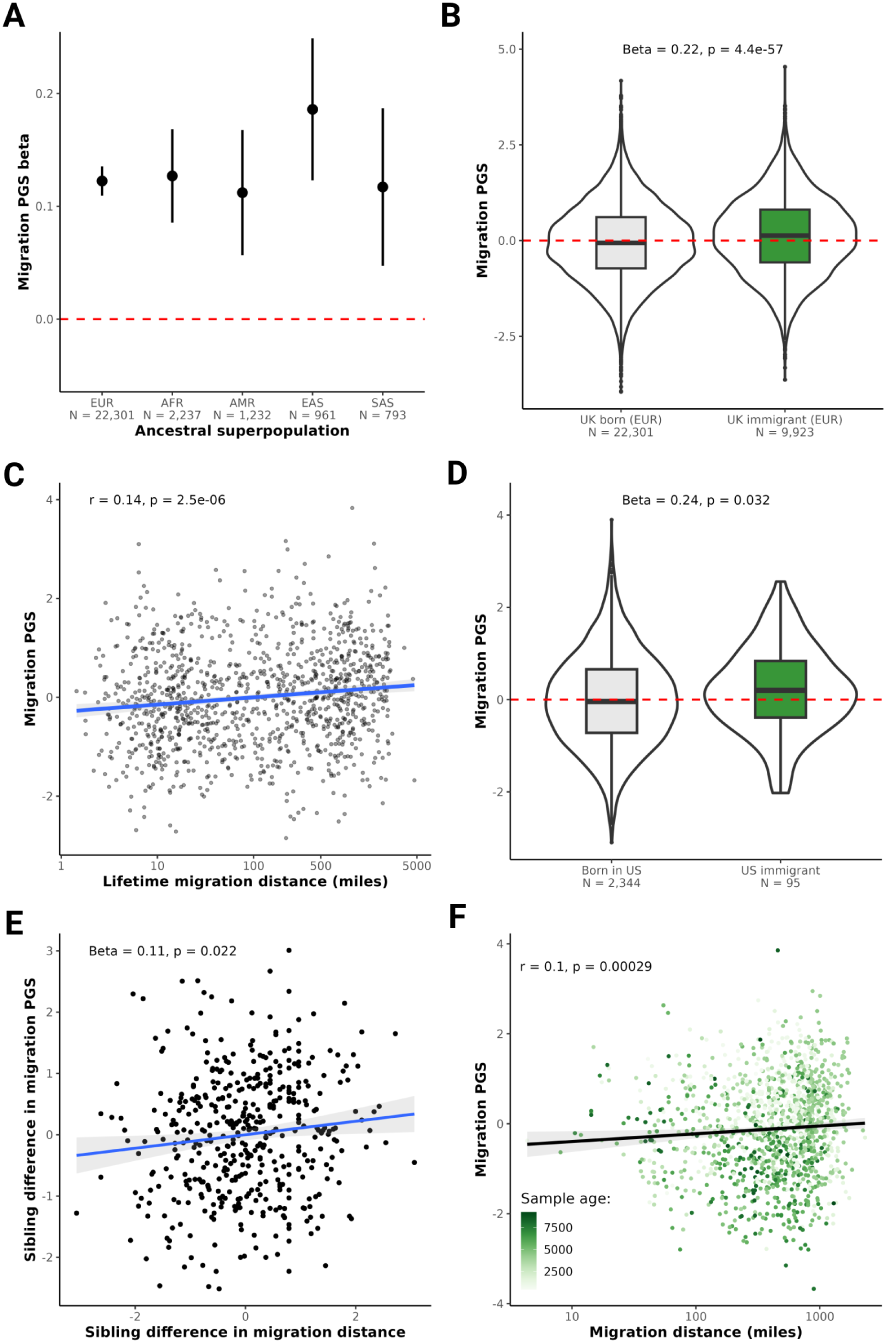
Polygenic predictions of migration generalize across populations and time. (**A**) Associations between the migration polygenic score (PGS) and actual migration distance within the UK across ancestral populations in the UK Biobank. Points represents *β* coefficients and 95% confidence intervals for each ancestral superpopulation. EUR = European (no overlap with discovery sample), AFR = African, AMR = American, EAS = East Asian, and SAS = South Asian. (**B**) Comparison of migration PGS between individuals born in the UK and still living there (grey) to UK immigrants (green). Logistic regression statistics are shown. (**C**) Correlation between lifetime migration distance in our US sample and the migration PGS (N = 1,128). Correlation statistics are shown. (**D**) Comparison of migration PGS between individuals born in the US and still living there (grey) to US immigrants (green). Logistic regression statistics are shown. (**E**) Sibling analysis results comparing the difference between each pair of siblings’ migration PGS and their difference in migration distance. Each point represents one sibling pair (N = 458 sibling pairs). Linear regression statistics are shown. (**F**) Correlation between inferred migration distance in ancient human samples from_24_the AADR and the migration PGS (N = 1,307). Correlation statistics are shown.

In our independent US sample, the migration PGS significantly predicted lifetime cumulative migration distance (*r* = 0.14, *p* = 2.5 × 10^−6^, N = 1,128; Figure 5C), accounting for approximately 2% of phenotypic variance. This association remained significant after adjusting for age, sex, and genetic principal components (*β* = 0.12, *p* = 4.6 × 10^−5^). The migration PGS predicted migration distance within individuals with autism (*r* = 0.14, *p* = 3.1 × 10^−4^, N = 690) and without (*r* = 0.13, *p* = 0.006, N = 438). Based on this sample, we estimate a 1 standard deviation increase in the migration PGS is associated with migrating 71.75 miles (115.5 kilometers) further away from home over one’s lifetime in the US.

To address potential confounding by population stratification, familial environment, and indirect genetic effects, we conducted a within-family analysis comparing sibling differences in migration PGS to sibling differences in migration distance. In UK Biobank sibling pairs (N = 458 pairs excluded from the GWAS sample), the migration PGS significantly predicted within-sibling-pair differences in migration distance (*β* = 0.11, *p* = 0.022; Figure 5E), demonstrating that the genetic effect persists even after controlling for environmental and genetic factors shared between siblings. This within-family analysis accounted for sibling differences in age, sex, genetic principal components, and educational attainment (years of completed education), indicating that the genetic propensity for migration carries information independent of educational differences.

Beyond predicting migration distance, we tested whether the PGS could distinguish immigrants from native-born individuals. In the UK Biobank, individuals born outside the UK (N = 9,923) showed significantly higher migration PGS compared to UK-born individuals (N = 22,301; *β* = 0.22, *p* = 4.4 × 10^−57^; Figure 5B). Similarly, in the US sample, immigrants (N = 95) had higher migration PGS than US-born individuals (N = 2,344; *β* = 0.24, *p* = 0.032; Figure 5D). These findings suggest that genetic propensity for migration predicts not only the distance individuals move within a country but also the likelihood of international migration.

The migration PGS also predicted outcomes outside of geographic mobility, suggesting that individual traits may create social environments that facilitate migration. In the US cohort, both migration distance and the migration PGS were associated with LGBTQ+ identification (phenotypic: *β* = 0.20, *p* = 1.5 × 10^−4^; PGS: *β* = 0.11, *p* = 0.034). Consistent with these associations, at the GWAS summary statistics level we observed a significant genetic correlation between migration distance and non-heterosexual behavior (**r*_*g*_* = 0.27, *p* = 2 × 10^−5^). Although LGBTQ+ individuals may migrate farther in search of more supportive environments, the observed genetic correlation indicates that shared heritable influences may also contribute, revealing how genetic and environmental pathways jointly shape migration behavior.

The convergence of between-family and within-family analyses, combined with cross-population replication, provides evidence that common genetic variants associated with migration propensity involve biological pathways rather than being entirely explained by population structure or socioeconomic confounding. The within-family design is particularly powerful because it controls for many factors shared between siblings, including ancestry, family environment, parental socioeconomic status, and neighborhood effects. These results suggest that the biological mechanisms underlying migratory behavior are largely shared across contemporary human populations.

### Migration polygenic score predicts behavior in ancient populations

To test whether genetic influences on migration extend across evolutionary timescales, we evaluated our UK-derived migration PGS in ancient human samples. We obtained genotype data and migration distance estimates (calculated from inferred birthplace to burial location) for 1,307 ancient individuals spanning approximately 500 to 10,000 years ago across West Eurasia (see Methods (*18, 19*)). The migration PGS significantly predicted migration distance in these ancient humans (*r* = 0.1, *p* = 2.9 × 10^−4^; Figure 5F), accounting for ≈1% of phenotypic variance. The prediction of migration behavior in ancient individuals demonstrates that the genetic architecture underlying human migration has been stable across evolutionary timescales.

### Migration alleles show evidence of recent positive selection

We examined whether migration-increasing alleles have systematically increased or decreased in frequency over time, which would indicate directional selection on migratory behavior. We calculated a selection coefficient by regressing migration PGS values against sample age across 3,241 ancient individuals spanning approximately 14,000 years. Migration-increasing alleles showed a significant increase in frequency over time (selection coefficient = 0.089, *p* = 3.2 × 10^−11^; Figure S2), indicating positive selection favoring migration propensity. Based on the observed selection coefficient, we estimate an increase of 0.101 standard deviations in migration PGS per 1,000 years. This finding was robust across analytical approaches. Genetic correlation analysis of allele frequency changes over the past 14,000 years confirmed that variants under positive selection (i.e., those increasing in frequency) are associated with greater migration distance (*r_*g*_* = 0.18, *p* = 9.6 × 10^−7^). To confirm that the genome-wide selection signal reflected selection on migration-associated variants rather than broader demographic processes, we examined directional selection specifically at our independent GWAS loci (N = 17 of 20 independent SNPs with selection information) (*36*). This locus-specific analysis demonstrated that effect alleles associated with greater migration distance have become significantly more common over time (*p* = 0.015; Figure S4), indicating directional concordance between allele frequency changes and phenotypic effects at these specific variants. For example, rs11710798, which was associated with further migration in the UK Biobank (*β* = 0.023, *p* = 1.7×10^−8^), has increased in frequency by ≈ 1% over the past 14,000 years, changing from 9% to 10% MAF. Conversely, rs73195301, which was associated with less migration distance (*β* = -0.016, *p* = 3.9 × 10^−8^), has decreased by ≈ 0.5%, from 23.2% to 22.8% MAF.

Next, to examine selection over deeper evolutionary time, we compared migration PGS across Neanderthal and Denisovan genomes (N = 10, mean age = 62,270 years, mean PGS = -2.97), ancient Europeans (N = 3,241, mean age = 3,217 years, mean PGS = -0.53), and modern Europeans (N = 503 from 1000 Genomes Project, mean PGS = 0). This analysis revealed a consistent pattern of increasing migration PGS from archaic hominins through ancient to modern humans (Supplementary Figure S3), suggesting sustained directional selection on migration-related alleles throughout recent human evolution. These findings indicate that human migration propensity has been under recent positive selection, with genetic variants promoting greater mobility systematically increasing in frequency. This selection pressure may reflect adaptive advantages of migration during periods of population expansion, environmental change, and agricultural transitions, suggesting that migratory behavior has been an evolutionarily advantageous strategy in recent human history.

### Exploratory analysis links migration-associated genetic variation to regional economic growth

Given our observed individual-level associations between migration genetics and socioeconomic outcomes, we tested whether population-level shifts in migration-associated alleles correlate with regional economic patterns. Across 222 US counties, we compared median PGS of current residents to those born locally to estimate county-level changes in PGS and examined their associations with per-capita income growth. This exploratory analysis has important limitations: our US sample was ascertained for autism research (70.2% ASD, 52.3% LGBTQ+), limiting population representativeness, and the observational design cannot establish whether genetic variation influences regional prosperity or economically dynamic regions selectively attract specific migrants. As a positive control, county-level education and income PGS significantly correlated with median income per capita (*r* = 0.25, *p* = 0.00014 and *r* = 0.21, *p* = 0.0018, respectively; Supplementary Figure S7). Counties experiencing increases in migration PGS showed significantly greater income growth per capita (*β* = 0.35, *p* = 0.0056; Figure 6B), as did counties with increasing income PGS (*β* = 0.3, *p* = 0.014) and education PGS (*β* = 0.24, *p* = 0.033; Figure 6). No significant associations were observed for other tested polygenic scores (Figure 6A). The migration PGS effect remained significant after controlling for changes in genetic principal components and baseline PGS (*β* = $3,855, *p* = 0.02). Based on these results, a 1 standard deviation increase in county-level migration PGS associates with $4,170 greater income growth per capita (95% CI: $1,246-$7,094), though this estimate carries substantial uncertainty given sample size and potential confounding. After additionally controlling for education PGS changes and genetic PC changes, the estimated effect decreases to $3,128 (*p* = 0.04, 95% CI: $102-$6,154), suggesting migration genetics capture macroeconomic associations partially independent of educational effects.

**Figure 6:**
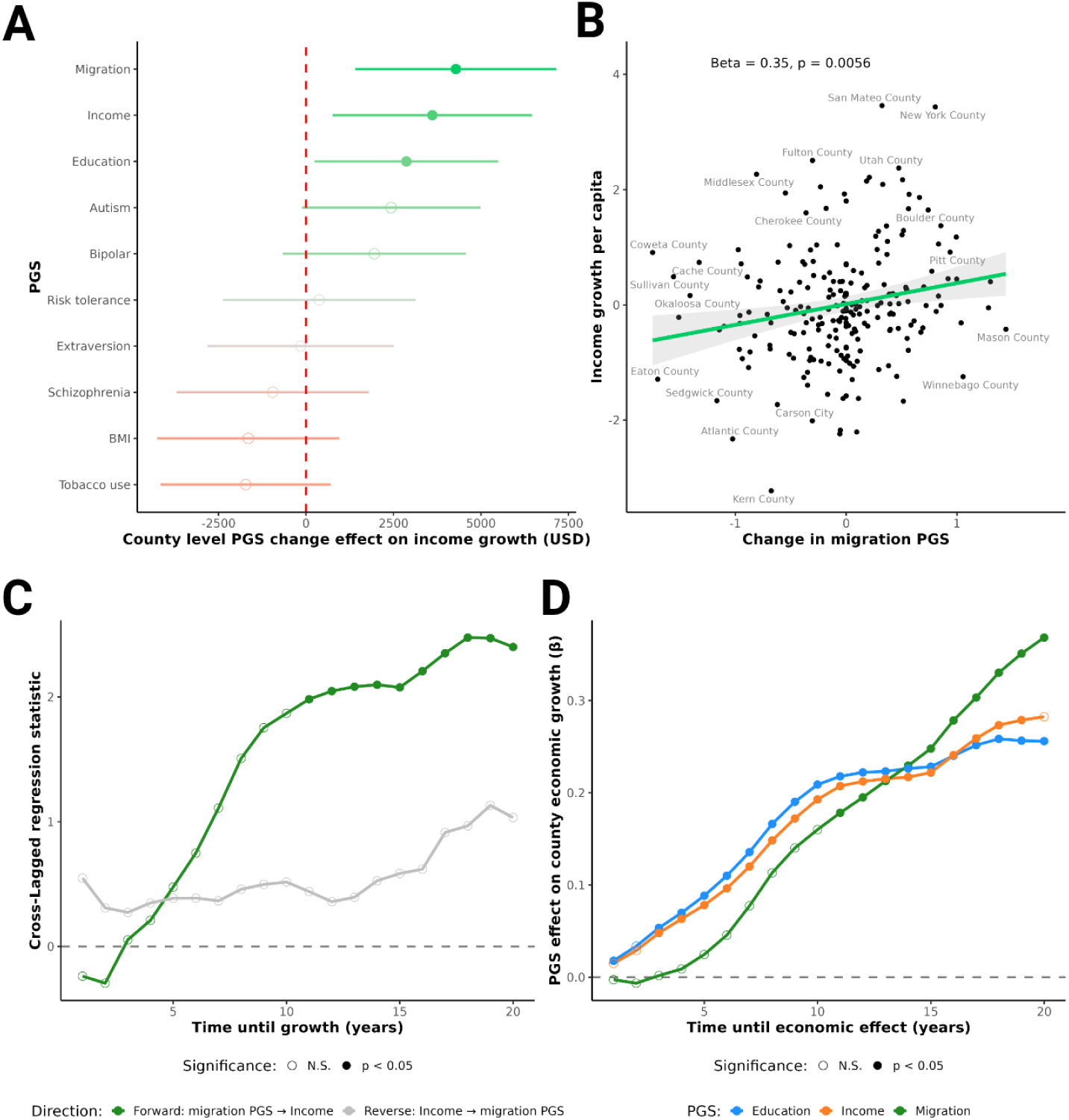
Migration PGS predicts economic growth across US counties. (**A**) Estimated effects of county-level changes in PGS on per-capita income growth (in USD). Points represent regression estimates and horizontal lines indicate 95% confidence intervals. Green indicates positive associations (e.g., counties where the median PGS increased over time had an increased in economic growth per capita over that same period). (**B**) Scatterplot illustrating relationship between income growth and county-level changes in migration PGS. Each point represents one county, with the regression line and 95% confidence interval shaded. Reported *β* and p-values are from linear regression estimates. (**C**) Cross-lagged panel analysis of migration PGS and county-level income growth. Green line shows forward effects (migration PGS at time (*t*) predicting future income growth at time (*t* + *ℓ*), controlling for baseline income); gray line shows reverse effects (income at time *t* predicting migration PGS change by time (*t* + *ℓ*), controlling for baseline PGS). Y-axis is the cross-lagged regression statistic. (**D**) Comparison of temporal dynamics of polygenic score changes predicting future economic growth for three polygenic scores (PGS): education PGS (blue), income PGS (orange), and migration PGS (green). 25

To distinguish whether genetic composition precedes economic growth or vice versa, we performed cross-lagged panel analyses testing both causal directions across lag intervals from 1 to 20 years (Figure 6C). The forward model examined whether county-level PGS at time *t* predicted income growth by time *t* + *ℓ*, controlling for baseline income. The reverse model tested whether baseline income predicted future changes in genetic composition via selective migration, controlling for baseline PGS. Results strongly supported a unidirectional relationship (Figure 6C). Increases in migration PGS predicted subsequent county economic growth, with effect sizes increasing consistently from near-zero at 1-year lags to *t* = 2.4 at 20-year lags (all *p <* 0.05 for lags ≥ 11 years). In contrast, the reverse pathway showed minimal evidence: income weakly predicted future PGS composition only at the longest time horizons (*t* = 1.13 at 19 years) and remained non-significant through all lags. This asymmetry indicates that genetic sorting drives regional economic divergence rather than simply reflecting migration responses to existing prosperity.

Comparison across polygenic indices revealed different temporal signatures (Figure 6D). Education PGS and income PGS predicting future economic growth almost immediately, with significant effects emerging within 1-3 years and growing steadily to *β* = 0.26-0.28 by 20 years. Migration PGS exhibited an incubation period before reaching significance, followed by rapid acceleration to the strongest long-term effects (*β* = 0.33 at 19 years, *p* = 0.014). This divergence suggests distinct mechanisms: education and income PGS capture direct human capital contributions upon arrival, whereas migration PGS operates through longer-term processes—perhaps community-building, entrepreneurship, or cultural dynamism—that compound over time to produce larger returns. Migration PGS ultimately showed the strongest effects on long-term economic growth in our sample, indicating that genetic propensity for migration may be an overlooked factor underlying regional economic differences.

## Discussion

We demonstrate that human migration is a heritable behavior shaped by genetic variants influencing gene expression in cortical excitatory neurons, with stable genetic architecture across at least 10,000 years of human history and evidence of recent positive selection. Stability and selection are not contradictory: the genetic architecture—which variants influence migration—has remained constant, while positive selection has increased the frequency of migration-promoting alleles in the population. Three independent analytical approaches (functional genomics, within-family analysis, and GWAS-by-subtraction) confirm that these genetic effects remain significant after accounting for gene-environment correlations arising from social stratification, consistent with biological influences on migration. Temporal increases in migration-associated genetic variants at the county level also correlate with regional economic growth in the contemporary United States. These findings bridge molecular neuroscience, evolutionary biology, and economics, revealing how heritable variation in complex behaviors can propagate from individual neurodevelopment to population-level outcomes across evolutionary and contemporary timescales.

Our GWAS identified 20 independent loci in established neurodevelopmental genes, including *CAMKV*, *MEF2C*, *BTN3A2*, *PCDH17*, and *TCF4* (*21, 23–25, 37*), with SNP-based heritability of approximately 5%. This places migration in the same heritability range as other complex behavioral traits such as risk-taking, extraversion, and neuroticism (*38, 39*). Extending recent work demonstrating genetic associations with migration distance (*11*), we incorporate functional genomics, GWAS-by-subtraction, and ancient DNA analyses to distinguish biological mechanisms from gene-environment correlations. While our findings validate previous reports of genetic correlations between migration and educational attainment and health outcomes (*12, 13, 40*), we provide several lines of evidence suggesting that migration genetics function through biological pathways rather than being fully explained by social stratification. Within-family analyses in 458 sibling pairs demonstrated that genetic effects persist even after controlling for shared environmental and genetic factors between siblings (*β* = 0.11), with effects remaining significant after adjusting for differences in education levels. This within-family design is particularly powerful because it controls for factors shared between siblings, including ancestry, family environment, parental socioeconomic status, and neighborhood effects. These are precisely the gene-environment correlations that can create spurious associations in between-family GWAS (*14*). Additionally, GWAS-by-subtraction analyses (*27*) confirmed significant independent heritability (SNP *h*^2^ = 1.9%) and identified a novel locus in *HOMER2*, establishing that migration captures genetic variation beyond educational attainment. The convergence of between-family, within-family, and GWAS-by-subtraction approaches establishes that migration propensity captures distinct motivational or exploratory tendencies not fully explained by education alone.

Functional genomic analyses revealed that migration-associated variants are most enriched in genes highly expressed in cortical excitatory neurons, particularly in layers 2/3, 5, and 6. These regions are critical for higher-order cognitive processing, decision-making, and long-range cortical projections (*41*). Migration alleles upregulate excitatory neuron marker genes at nearly four times the rate of inhibitory neuron markers (odds ratio = 3.67), pointing to systematic effects on excitatory neuron biology. The enrichment in prenatal brain tissue further suggests that migration propensity emerges through early neurodevelopmental processes that shape cortical architecture. This cellular mechanism links migration genetics to evolutionarily expanded neural substrates underlying cognitive abilities that facilitate environmental adaptation and risk-taking (*42, 43*). Excitatory neurons in these cortical layers exhibit increased dendritic complexity, larger cell sizes, and greater synaptic integration in humans compared to other species (*43*), with these cellular properties correlating with individual differences in intelligence and cognitive flexibility (*44*). This body of evidence explains why migration propensity strongly associates with education, cognition, openness, risk tolerance, and reduced interpersonal attachment. Taken together, the genetic, cellular, and cognitive evidence indicates that migration represents a complex behavioral output of neurodevelopmental processes shaping cortical excitatory function and cognitive flexibility.

The genetic architecture of migration is ancient. Our polygenic score, derived from modern UK residents, predicted inferred mobility in 1,307 individuals who lived up to 10,000 years ago, from Neolithic farmers to Bronze Age nomads (*r* = 0.1). Showing the migration polygenic score predicts behavior in people who died thousands of years ago demonstrates remarkable stability of genetic architecture despite profound changes in technology, social organization, and environmental contexts. The same PGS predicted migration distance across all five ancestral superpopulations in the UK Biobank with consistent effect sizes (*β* = 0.11 to 0.16), replicated in an independent US cohort (*r* = 0.14), and distinguished immigrants from native-born individuals in both countries. Together, these results suggest the same biological mechanisms (exploratory drive, risk tolerance, and cognitive flexibility) have likely driven human mobility across fundamentally different cultural contexts, from hunter-gatherer range expansions to modern intercontinental relocations.

Ancient genomes enabled direct tests of natural selection, revealing robust evidence of positive selection on migration-associated alleles. The average migration PGS increased by approximately 1 standard deviation over the past 10,000 years. Migration polygenic scores increased progressively from Neanderthals through ancient to modern humans, consistent with sustained directional selection throughout recent human evolution. The ability to relocate and establish communities in new environments likely conferred substantial fitness benefits during periods of population expansion, agricultural transitions, and environmental change. The correspondence between migration allele frequency increases and major cultural and linguistic transitions adds to growing evidence that polygenic adaptation on behavioral and cognitive traits has been pervasive during the past 10,000 years (*36*).

Cross-lagged panel analysis supported a directional relationship: counties that accumulated more migration-associated genetic variation predicted subsequent income growth—roughly $3,100 more per person per standard deviation increase in county-level migration PGS, even after controlling for education-related genetic differences (95% CI: $102-$6,154)—while the reverse pathway (income predicting genetic composition) showed minimal evidence. Important caveats remain: our US sample was ascertained for autism research with LGBTQ+ enrichment, limiting population representativeness, and unmeasured demographic factors may confound these associations. Replication in representative samples is needed, but the cross-lagged evidence suggests that individual-level associations between migration genetic propensity may aggregate to regional effects through selective migration and human capital concentration.

Our findings weigh evidence across four major hypotheses for individual migration propensity (Table 1) and show that genetic influences play a modest but significant role. Environmental determinism cannot account for the observed heritability (h^2^ = 5%) or within-family differences (*β* = 0.11). Population-stratification explanations are contradicted by multiple independent lines of evidence: within-family effects persist; migration variants are functionally enriched in cortical excitatory neurons; GWAS-by-subtraction isolates variation independent of education; the PGS replicates across ancestries and predicts behavior in ancient populations; and migration alleles show signatures of positive selection. Instead, the results most strongly support heritable individual-difference models, suggesting that neurodevelopmental pathways shaping cognitive flexibility and exploratory behavior contribute to migration propensity and have been evolutionarily stable over at least 10,000 years. At the same time, the data are consistent with gene–environment interplay: heritable traits likely influence how individuals perceive and respond to opportunities, but environmental conditions remain essential to realized migration outcomes.

**Table 1:** Core hypotheses for individual migration differences and support from this study.

| Hypothesis | Prediction | Support from Current Study |
| --- | --- | --- |
| **Environmental Determinism** | Migration driven entirely by external conditions (economic opportunities, conflict, etc.); individuals respond uniformly to circumstances; no systematic heritable individual differences. | **Not supported:** Cannot explain SNP $h^2 = 5\%$, within-family effects ($\beta = 0.11$, $p = 0.022$), or sibling discordance despite identical birthplaces and similar family environments. |
| **Population Stratification Artifact** | Apparent genetic associations arise from population stratification, assortative mating, and indirect genetic effects; effects disappear in within-family analyses; no functional biological mechanisms. | **Not supported:** Within-family effects persist after controlling for shared environment, birthplace, and education; functional enrichment in cortical excitatory neurons; GWAS-by-subtraction identifies independent loci from education; PGS predicts across five ancestries and in ancient populations where modern stratification cannot operate. |
| **Heritable Individual Differences** | Biologically-influenced traits shape migration propensity independent of shared environment; genetic effects persist within families; specific neural mechanisms identifiable. | **Supported:** Significant SNP $h^2 = 5\%$; within-family effects ($\beta = 0.11$) after controlling for shared environment and education; functional enrichment in prenatal brain and cortical excitatory neurons; PGS predicts ancient migration; positive selection over 10,000 years; consistent replication across five ancestries. |
| **Gene-Environment Interplay** | Heritable traits and environmental factors jointly determine migration; genetic propensities influence how individuals perceive and respond to migration opportunities. | **Supported:** Genetic effects evident (within-family, functional enrichment, ancient DNA) alongside strong environmental associations (phenotypic correlations with socioeconomic factors); consistent with heritable traits shaping responses to opportunities. |

In conclusion, human migration is a heritable behavior with deep evolutionary roots: shaped by genetic variants active in cortical excitatory neurons, with genetic architecture conserved across 10,000 years even as positive selection has increased the frequency of migration-promoting alleles. By integrating functional genomics, within-family designs, ancient DNA, and population-level economic data, we trace a causal chain from prenatal neurodevelopment through individual behavior to regional economic outcomes. These findings reframe migration as not merely a response to circumstance, but a behavioral strategy with biological underpinnings that natural selection has consistently favored. Understanding why some individuals are drawn to move, and what they bring when they do, has implications that extend from evolutionary biology to contemporary debates about human mobility.

## Materials and methods summary

We quantified migration distance as the Haversine distance between birthplace and current/final residence coordinates in two samples: (1) the UK Biobank (*16*) participants born in the UK, and (2) a sample of 3,235 US adults enriched for autism and LGBTQ+ identities (70.2% ASD and 52.3% identifying as LGBTQ+) recruited with the SPARK Research Match platform (*17, 45*) as part of a larger study investigating the relationship between mental health, stress, gender identity, and sexuality. We examined phenotypic correlations between migration distance and cognitive, personality, health, and socioeconomic measures using linear regression adjusted for relevant covariates (e.g., age and sex).

We conducted a genome-wide association study (GWAS) of log10-transformed migration distance in 242,561 unrelated individuals of White British ancestry from the UK Biobank using LDAK-KVIK (*46*), adjusting for age, sex, genetic principal components, and birthplace neighborhood socioeconomic status (Townsend deprivation index). SNP-based heritability and genetic correlations with 48 other traits were estimated using LD Score Regression (LDSC) (*26*). GWAS-by-subtraction (*27*) was conducted using Genomic Structural Equation Modeling (*47*) (implemented with the GSUB tool (*48*)) to identify migration-related genetic variation independent of educational attainment.

We mapped migration-associated variants from our GWAS to tissues and cell types using MAGMA (*34,35*), gsMap (*28*) (spatial transcriptomics in E16.5 mouse embryo (*29*)), and scPagwas (*30*) (human postmortem medial temporal cortex single-cell RNA-seq (*49*)). Cell-type-specific eQTL analysis correlated migration GWAS Z-scores with expression effects in excitatory and inhibitory neurons using the singleBrain meta-analysis with two sets of neuronal marker genes (*31, 32*).

Migration polygenic scores (PGS) were constructed using LDpred2-inf (*50*) for contemporary samples (UK Biobank and US samples) and PRSice-2 (*51*) with clumping/thresholding for ancient samples. We validated PGS predictions across five ancestral superpopulations in the UK Biobank (N=24,323), in an independent US cohort of 3,235 adults recruited from the SPARK Research Match platform (*17, 45*), within 458 UK Biobank sibling pairs, and for predicting immigration status in both UK and US samples.

To determine whether the effects observed in contemporary humans are relevant to ancient humans and whether these alleles may have been advantageous, we analyzed 3,241 ancient human genomes from the Allen Ancient DNA Resource (AADR v54 (*18*)) spanning 150-14,000 years before present across West Eurasia, including 1,307 individuals with inferred migration distances that have been previously validated (*19*). Our polygenic analysis regressed PGS on sample age using linear mixed models (accounting for population structure and relatedness), with validation via LDSC genetic correlation of allele frequency changes (*36*), as well as comparisons between archaic humans, ancient humans, and modern humans. We validated these results with a SNP level analysis of independent GWAS migration hits (N = 17 of 20 loci with selection information), testing for directional selection at the variant level.

For population-level economic analysis, we calculated median PGS for individuals born in versus currently residing in 222 US counties, testing associations between county-level PGS changes (based on our US sample enriched for autism and LGBTQ+ identity) and economic growth (2023 income residualized on 1980 income) using weighted linear regression (weighted by population size). To account for potential population stratification, we also ran models that included changes in genetic principal components as covariates. A follow-up analysis used these data for cross-lagged panel models, testing whether migration PGS predicts future economic growth or income predicts future PGS change. Economic and population size measures came from the Bureau for Economic Analysis.

## Supporting information

All supplementary tables

## General

We are grateful to all of the participants and families in the UK Biobank, SPARK, the SPARK research match team and Tempus, the SPARK clinical sites, and research staff. We appreciate obtaining access to genetic and phenotypic data for SPARK data on SFARI Base. We are incredibly appreciative of the open access datasets used in this study, including the researchers who put together the AADR dataset.

## Funding

This work was supported by the National Institutes of Health (HG012697 to JFS and JJM). This work was also supported by the Roy J. Carver Charitable Trust through a professorship to JJM. Additional support for this work came from the University of Iowa Hawkeye Intellectual and Developmental Disabilities Research Center (Hawk-IDDRC) through the Eunice Kennedy Shriver National Institute of Child Health and Human Development (P50HD103556), and the Simons Foundation (SFARI and SPARK) to JJM and LGC.

## Author contributions

The study was designed by LGC and JJM. The SPARK Research Match study was carried out by AT with guidance by JSY, JFS, and JJM. The data was processed by LGC, AT, ME, and JJM. The analyses were performed by LGC and JJM. The manuscript was written by all authors.

## Competing interests

There are no competing interests to declare.

## Data and materials availability

GWAS summary statistics can be downloaded here:

https://zenodo.org/records/18472458

UK Biobank data are available to approved researchers at:

https://www.ukbiobank.ac.uk/enable-your-research/apply-for-access

SPARK genetic and phenotype data is available to qualified researchers at SFARI base:

https://base.sfari.org/

Allen Ancient DNA Resource:

https://reich.hms.harvard.edu/allen-ancient-dna-resource-aadr-downloadable-genotypes-

We used publicly available tools for data processing and analysis: PLINK:

https://www.cog-genomics.org/plink

LDAK-KVIK:

https://www.ldak-kvik.com/

LDSC:

https://github.com/bulik/ldsc

FUMA:

https://fuma.ctglab.nl/

Genomic SEM:

https://github.com/GenomicSEM/GenomicSEM

GSUB:

https://github.com/qlu-lab/GSUB

GCTA:

https://yanglab.westlake.edu.cn/software/gcta

LDpred2:

https://privefl.github.io/bigsnpr/articles/LDpred2.html

PRSice-2:

https://choishingwan.github.io/PRSice/

R:

https://www.r-project.org/

## Supplementary Materials for

### Materials and Methods

#### Human subjects and IRB

UK Biobank analyses were conducted under application number 46053. US sample collection was approved by the University of Iowa IRB (201611784) and WCG (formerly known as WIRB; 201703201). All participants provided informed consent. Ancient DNA data were obtained from published studies with appropriate ethical approvals.

#### UK Biobank

The UK Biobank is a prospective cohort study of approximately 500,000 participants aged 40-69 years recruited between 2006-2010 (*16*). For phenotypic analyses, we included approximately 390,000 individuals of European ancestry with complete migration distance and covariate data. For genome-wide association analysis, we restricted to 242,561 unrelated individuals of White British ancestry with phenotypic and covariate data (genetic relatedness cutoff *<* 0.05). For cross-ancestry validation, we analyzed independent samples from five ancestral superpopulations: European (N = 19,409), African (N = 2,108), Admixed American (N = 1,163), East Asian (N = 904), and South Asian (N = 739). For within-family analysis, we identified 458 sibling pairs not included in the GWAS discovery sample using established kinship thresholds (IBS0 *>* 0.1% and kinship coefficient 0.18-0.35) (*52*), further restricted to pairs with identical reported birthplaces to control for differences in early environmental factors.

#### US sample

We recruited 3,235 adults (≥ 30 years old) from the SPARK cohort through the Research Match platform (*17, 45*) as part of a larger study investigating the relationship between mental health, stress, gender identity, and sexuality. As part of this study, participants provided detailed residential histories including all previous zip codes, enabling calculation of lifetime cumulative migration distance. The analytic sample included 1,128 individuals with complete genotype and residential history data for PGS validation analyses who reported at least 1 move, and 2,439 individuals for immigration status analyses (95 immigrants, 2,344 US-born). We limited the migration distance analysis to individuals from the contiguous United States, removing individuals who reported being born in or currently living in Hawaii or Alaska.

#### Ancient DNA

We obtained ancient genomes from the Allen Ancient DNA Resource (AADR v54 (*18*)). After quality control, our dataset comprised 3,241 ancient *Homo sapiens* individuals spanning 150-14,000 years before present across West Eurasia (longitude 25W-60E, latitude 35N-80N). Quality control filters included: AADR assessment status “PASS”, *<* 50% missingness at common autosomal SNPs, *<* 10% missingness per SNP, and removal of duplicate/twin pairs (genetic relatedness *>* 0.9). A subset of 1,307 individuals had inferred migration distances based on genetic ancestries (*19*). For deep-time comparisons, we also included 10 archaic hominin genomes from the AADR (Neanderthal and Denisovan) and 503 modern Europeans from the 1000 Genomes Project (*53*).

### Phenotype definitions

Migration distance was defined as the Haversine distance between birthplace and current/final residence coordinates. In the UK Biobank, this was calculated from reported birthplace coordinates to residential address at recruitment (approximated by assessment center location). In the US sample, lifetime cumulative migration distance was calculated as the sum of distances between all sequential residences. For ancient individuals, migration distance was the distance from inferred birthplace to burial location (*19*). All migration distances were log10-transformed (+1 mile) to handle zero values and approximate normality. Results were robust to rank inverse normal transformation.

### Genotyping and quality control

Genotypes for UK Biobank participants were obtained from the v3 imputation release. Briefly, participants were genotyped using the UK BiLEVE Axiom or UK Biobank Axiom arrays and imputed to a custom reference panel combining the Haplotype Reference Consortium (HRC), UK10K, and 1000 Genomes samples. Genotype quality control included: sample missingness *<* 2%, SNP missingness *<* 5%, imputation quality (INFO) *>* 0.6, and minor allele frequency (MAF) *>* 0.5%. The US sample (SPARK) underwent similar quality control but were imputed to the TopMed reference panel (*54*) to capture greater genetic diversity, with more stringent SNP filtering: missingness *<* 2% and INFO *>* 0.8. Ancient DNA genotypes were obtained from the curated AADR published dataset (version 54 (*18*)) and used as available without additional imputation, given the challenges of imputing ancient genomes.

### Genome-wide association study

We performed GWAS of log10-transformed migration distance in 242,561 unrelated White British individuals using LDAK-KVIK (*46*). We opted to use the LDAK-KVIK method because it adjusts for false positives, population structure, and cryptic relatedness better than standard regression approaches and offers improved power over other mixed-model approaches. For calculating the leave-one-chromosome-out polygenic prediction component of the LDAK-KVIK pipeline, we used LD pruned HapMap3+ SNPs (*55*) (pruned using PLINK (*56*) with the command “–indep-pairwise 1000 100 0.9”). The phenotype was residualized for age, age^2^, sex, first 20 genetic principal components, retirement status, UK country of birth, and birthplace Townsend deprivation index prior to analysis. Lead SNPs at genome-wide significant loci (p *<* 5 × 10^−8^) were annotated to the nearest gene based on hg19 coordinates. SNP-based heritability 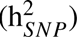 was estimated using LD Score Regression (LDSC) with the 1000 Genomes European reference panel (*26, 53*).

To identify approximately independent genome-wide significant loci, we performed linkage disequilibrium (LD) clumping using PLINK2 (*57*). Clumping was conducted with European ancestry reference data from the 1000 Genomes Project (*53*). LD clumps were defined using a primary and secondary significance threshold of P *<* 5 × 10^−8^, an LD threshold of *R*^2^ *<* 0.1, and a ±1Mb window around each index variant.

### Genetic correlation analysis

Genetic correlations (r*_*g*_*) between migration distance and published GWAS of cognition (cognitive performance (*58*), executive functioning (*59*), brain volume (*60*)), socioeconomic related (income (*61*), education (*62*), Townsend Deprivation index (*63*), partner choice index (*64*), population density (*65*)), neurodevelopmental and psychiatric conditions (autism (*66*), bipolar (*67*), anorexia (*68*), OCD (*69*), Tourette’s (*70*), major depression (*71*), Schizophrenia (*72*), addiction (*73*), anxiety, insomnia, cross disorder, ADHD (*74*), PTSD (*75*), Alzheimer’s (*76*), suicide attempt), personality (risky taking behavior (*38*), loneliness, friend satisfaction, family satisfaction, work satisfaction, financial satisfaction, BIG5 personality traits (*39*)), behavioral (externalizing problems (*77*), aggression (*78*), antisocial behavior (*79*), age at first birth, age at first sexual intercourse, regular gym attendance, regular religious service attendance, regular pub attendance), and health traits (self-reported health, asthma, COPD, BMI, longevity, birth weight (*80*)) were estimated using LDSC (*26*). Summary statistics underwent standard quality control (INFO *>* 0.6, MAF *>* 0.01, restriction to HapMap3 SNPs). To account for possible inflation of test statistics due to population stratification, we constrained the LDSC intercept prior to genetic correlation calculation. Multiple testing correction used the false discovery rate (FDR) method.

### GWAS-by-subtraction

To identify migration-related genetic variation shared and independent of educational attainment, we performed GWAS-by-subtraction using Genomic Structural Equation Modeling (Genomic SEM) (*47*) implemented in the GSUB package (*48*) using the 1000 Genomes European reference panel and a MAF filter of 1%. Full genetic correlations for each factor can be found in Supplemental tables 8. Genome-wide significant independent loci can be found in Supplemental tables 6-7. GWAS results of the shared migration-education factor (we call this the “Major” factor) and the component of migration independent of education (“Minor” factor) can be seen in Figure S1.

### Functional genomics

#### Tissue and spatial enrichment

We mapped migration-associated variants to tissues using gsMap (*28*), which integrates GWAS summary statistics with spatial transcriptomics. Analysis used mouse embryo data from the E16.5 timepoint (*29*) with default parameters to identify regions of enriched heritability for migration. Results were validated using MAGMA (*34*) gene-set enrichment analysis with GTEx V8 tissues (*81*) through the FUMA platform (*82*), applying FDR correction. gsMap results can be found in Supplemental table 14. MAGMA results for GTEx v8 tissues can be found in Supplemental Tables 11-12.

#### Cell-type enrichment

Cell-type-specific heritability enrichment was assessed using scPagwas (*30*) applied to the SEA-AD single-cell RNA-sequencing atlas of human postmortem medial temporal cortex (*49*). We tested enrichment across major cell types (excitatory neurons, inhibitory neurons, astrocytes, oligodendrocytes, microglia, endothelial cells) and neuronal subtypes defined by cortical layer and projection pattern. Results were validated using MAGMA (*34,35*) cell-type specific gene-set enrichment analysis through the FUMA (*82*) platform with PsychENCODE developmental brain cell-type specific data (*83*) and a large single-cell prenatal brain atlas (*84*) (we focus on results from 16 weeks post-conception, the timepoint with the strongest associations, across all provided brain regions), applying FDR correction. scPagwas results can be found in Supplemental table 15. MAGMA cell-type enrichment results can be found in Supplemental Table 13.

#### Cell-type-specific directional eQTL analysis

To determine directional effects of migration variants on cell-type-specific gene expression, we correlated migration GWAS Z-scores with single-cell eQTL Z-scores from excitatory and inhibitory neurons derived from the singleBrain meta-analysis (*32*).

For the primary analysis, cell-type-specific marker genes for excitatory (glutamatergic) and inhibitory (GABAergic) neurons were identified from the DESCARTES human fetal gene expression atlas (*31*). High-confidence markers were defined using stringent criteria: genes with expression *>*100 TPM in the marker cell type, fold-change *>*1.5 relative to comparator cell types (astrocytes, oligodendrocytes, microglia, and the opposite neuronal subtype), and differential expression *q <* 0.05. All markers were restricted to protein-coding genes present in the MAGMA gene analysis set (genes with sufficient proximal SNPs analyzed in the migration GWAS).

For each marker gene, we computed the correlation between migration GWAS effect sizes (Z-scores) and cell-type-specific eQTL effect sizes (Z-scores) for excitatory and inhibitory neurons separately. Positive correlations indicate that migration-increasing alleles increase gene expression in that cell type. We tested whether excitatory markers showed systematically different migration-eQTL correlations than inhibitory markers using weighted logistic regression. Weights were defined as log_2_(TPM) to account for differential marker gene expression levels.

As a validation analysis, we repeated this procedure using an independent marker gene set from PanglaoDB (filtered to human marker genes labeled as markers for “Neurons”, “Glutaminergic neurons”, or “GABAergic neurons”) (*33*). For each gene, positive correlation indicates migration-increasing alleles increase expression. We classified genes as showing excitatory-specific effects (migration alleles increase expression in excitatory but decrease in inhibitory neurons), inhibitory-specific effects (opposite pattern), or non-specific effects (shared directional effects). Enrichment of excitatory-specific versus inhibitory-specific effects was tested using a proportion test, and an odds ratio was calculated based on the total number of subtype-specific genes. Gene-level celltype-specific associations between eQTLs and the migration GWAS are reported in Supplementary Table 16.

### Polygenic score construction

Migration polygenic scores were constructed using LDpred2-inf (infinitesimal model (*55*)) from UK Biobank GWAS summary statistics and applied to held-out UK Biobank participants and the US SPARK sample. More information on SPARK genotype processing and imputation can be found here (*85*). We used HapMap3+ SNPs with UK Biobank LD reference data provided as part of the LDpred2 package. Scores were residualized for the first 20 genetic principal components and standardized (mean = 0, SD = 1) within each analysis sample. For ancient samples, we used PRSice-2 (*51*) with clumping and thresholding (LD-independent SNPs, MAF *>* 1%, p-value threshold = 1) due to likely differences in LD structure, with standardization relative to 1000 Genomes Europeans (*53*).

### Polygenic score validation

#### Cross-ancestry validation

We tested migration PGS associations with log10-transformed migration distance across five ancestral superpopulations in the UK Biobank using linear regression, adjusting for age, sex, and principal components (PCs 1-20). Summary statistics for the association between the migration PGS and actual migration distance for each ancestral group can be found in Supplemental Table 10.

#### Ancestry assignment for cross-ancestry validation

We assigned UK Biobank participants to ancestral superpopulations using an integrated approach combining genetic clustering and self-reported ethnicity, following established protocols (*86*). We merged held-out UK Biobank samples (not included in GWAS discovery or related to anyone in the GWAS sample) with the 1000 Genomes Project reference panel (*53*) and performed principal component analysis on UK Biobank samples, projecting 1000 Genomes individuals onto the same PC space. We applied k-means clustering (k = 5) to genetic PCs in the 1000 Genomes samples to define cluster centroids for each superpopulation (EUR, AFR, AMR, EAS, SAS). UK Biobank participants were assigned to the nearest cluster based on Euclidean distance to cluster centroids. Final superpopulation assignments integrated genetic clustering with self-reported ethnicity: for example, individuals reporting “White” or “Irish” background and/or clustering with 1000 Genomes Europeans were assigned to the EUR superpopulation.

#### Replication in independent cohort enriched for autism and LGBTQ+ identities

In the US sample (70.2% with autism and 52.3% identifying as LGBTQ+), we tested PGS associations (calculated with LDpred2-inf (*55*)) with cumulative lifetime migration distance using correlation and linear regression with covariates (age, *age*^2^, sex, PCs 1-5). Variance explained (R^2^) was calculated as the squared correlation coefficient between observed and PGS-predicted phenotypes.

To explore how other traits may influence migration, we used logistic regression models to predict self-reported LGBTQ+ identity from both the migration PGS and actual phenotypic migration distance, controlling for age, *age*^2^, sex, and the first five genetic principal components. After adjusting for ASD status in our US sample, the observed association between the migration PGS and LGBTQ+ identification was no longer statistically significant, though the effect direction remained consistent (*β* = 0.08, *p* = 0.16). In contrast, actual migration distance remained nominally associated with LGBTQ+ identification (*β* = 0.12, *p* = 0.037). Given the correlational nature of this analysis and limited sample size, causal inference is not possible. However, these results may suggest that among individuals with ASD, components of the migration PGS may share genetic architecture with biological factors contributing to the elevated prevalence of LGBTQ+ identities observed in this population (*87–89*).

#### Within-family validation

For each of 458 sibling pairs, we calculated within-pair differences in migration distance, migration PGS, and all covariates (age, sex, year of birth, educational attainment, PCs 1-5). Migration distance differences were rank inverse normal transformed. We tested association between PGS differences and migration distance differences using linear regression, with and without covariate adjustment. For robustness, we also did a median split analysis and found sibling pairs that migrated further away from each other had larger differences in the migration PGS (*β* = 0.25, *p* = 0.009; S8).

#### Immigration status prediction

We compared migration PGS between immigrants and native-born individuals using logistic regression (covariates: age, sex, PCs 1-20 in UK and PCs 1-5 in US). Analyses were conducted separately in UK Biobank (UK-born: N = 19,409; immigrants: N = 9,923, all in the EUR superpopulation) and US sample (US-born: N = 2,344; immigrants: N = 95, mixed ancestral backgrounds).

### Ancient DNA analyses

#### Phenotypic prediction

We tested whether migration PGS predicted inferred migration distance in 1,307 ancient individuals using Spearman correlations (*19*). To ensure robustness, we analyzed two estimates of inferred migration distances (a PCA-based measure and an MDS-based measure). Both measures yielded concordant results; we report the PCA-based measure in the main text (*r* = 0.1 vs. *r* = 0.085 for MDS) because it incorporated more genomic information (5 principal components vs. 2 MDS dimensions).

#### Selection analysis

To test for directional selection, we regressed migration PGS on log10 transformed sample age (years before present) across 3,241 ancient individuals using linear mixed models to account for population structure and genetic relatedness, following recommended protocols (*36*). We also calculated the selection coefficient as the standardized change in PGS per 1,000 years. As validation, we performed genetic correlation analysis using LDSC to test whether alleles increasing in frequency over time were associated with higher migration distance (*36*). As further validation, we examined whether independent genome-wide significant migration loci were under directional selection. We overlapped the 20 independent migration loci with selection summary statistics from the AADR (N = 17 / 20 SNPs overlapping) (*36*). To test for directional selection at significant migration loci, we grouped the 17 overlapping SNPs by the direction of their migration association (positive or negative) and used Wilcoxon tests to compare allele frequency changes over the past 14,000 years between groups. SNP level data from this analysis can be found in Supplemental Table 9. All three groups were scored using identical SNP sets. For the deeper evolutionary time analysis, we compared mean PGS across archaic hominins, ancient Europeans, and modern Europeans using t-tests.

### County-level economic analyses

For 222 US counties with ≥ 2 participants born in the county and ≥ 2 current residents, we calculated median PGS for individuals born in each county and median PGS for current residents. Individuals reported birth and current zip codes were mapped to the nearest county, if an individual was *>* 40 miles away from the center of the county they were excluded from the analysis. County-level PGS change was defined as the difference between current resident PGS and birthplace PGS. County level income per capita data (1980 and 2023) were downloaded from the Bureau of Economic Analysis. Economic growth was defined as 2023 income residualized on 1980 income (1980 was the average birth year of our US sample).

We first validated the county-level approach by testing correlations between county-level education/income PGS and median income per capita using Pearson correlation. We then tested associations between county-level PGS changes (migration, education, income, autism, bipolar disorder, risk tolerance, extraversion, schizophrenia, BMI, tobacco use) and economic growth using linear regression. Linear regression models were weighted based on log10 transformations of population size in 2023 reported from the Bureau of Economic Analysis. Robustness analyses adjusted for changes in the first 5 genetic principal components (difference in median PC values between birthplace and current resident populations).

Next, to examine bidirectional temporal relationships between county-level polygenic scores (PGS) and economic outcomes we used a cross-lagged panel design. We computed PGS differences between current and historical average PGS values for each county (reflecting genetic composition changes due to selective migration) and median household income per capita log-transformed and standardized within each year. For each lag interval *ℓ* (1–20 years), we constructed two complementary models to test alternative causal pathways. The forward model tested whether baseline PGS predicts subsequent income growth, controlling for baseline income:

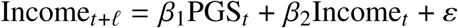

The reverse model assessed whether income predicts future genetic composition changes (via selective migration), controlling for PGS:

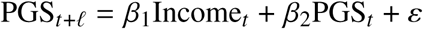

Models were estimated using ordinary least squares linear regression with cluster-robust standard errors (clustered by county) to account for within-county correlation over time. Analyses included only county-year observations with complete data for all variables. To compare the relative predictive power of across different polygenic scores, we estimated separate forward models for three polygenic scores: migration PGS, educational attainment PGS (*90*), and income PGS (*61*). Each model followed the same forward specification above, with PGS replaced by the respective polygenic score while controlling for baseline income.

### Statistical analysis

All analyses were performed in R version 4.3.1 (*91*). Additional software: PLINK 1.9/2.0 (*56, 92*), GCTA (*93*), LDSC (*26*), LDAK-KVIK, PRSice-2 (*51*), LDpred2 (*50*), FUMA (*82*), MAGMA (*34*), Genomic SEM (*47*), and GSUB (*48*).

## Supplementary Figures

**Figure S1:**
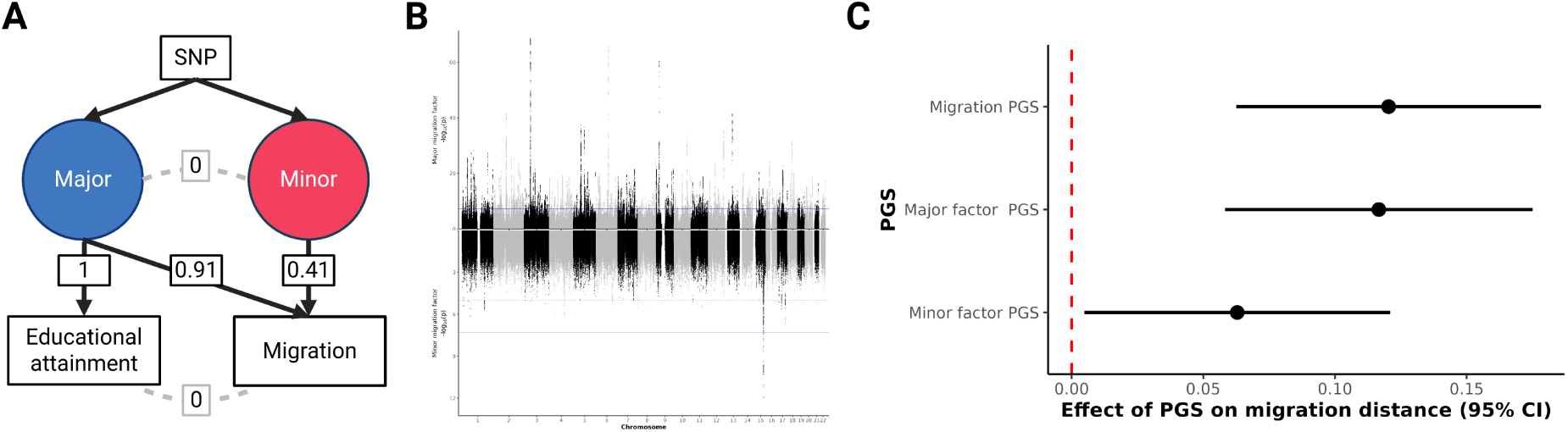
GWAS-by-subtraction reveals shared and distinct factors underlying migration and education. **A** Genomic SEM model fit. Major and Minor factors are latent (unobserved) variables estimated based on the GWAS summary statistics of migration and educational attainment. The covariances between migration and educational attainment and between the Major and Minor factors are fixed to 0. **B** Top panel of the Miami plot shows the GWAS results for the Major migration factor (the genetic component of migration shared with education). The bottom panel represents the Minor factor (the genetic component of migration that is distinct from education). Blue line indicates genome-wide significance (*p <* 5 × 10^−8^). **C** Association between lifetime migration distance in our US sample and the different migration PGS (N = 1,128). Regression *β* estimates and 95% confidence intervals are shown. Regression models included age, *age*^2^, sex, and 5 genetic PCs as covariates.

**Figure S2:**
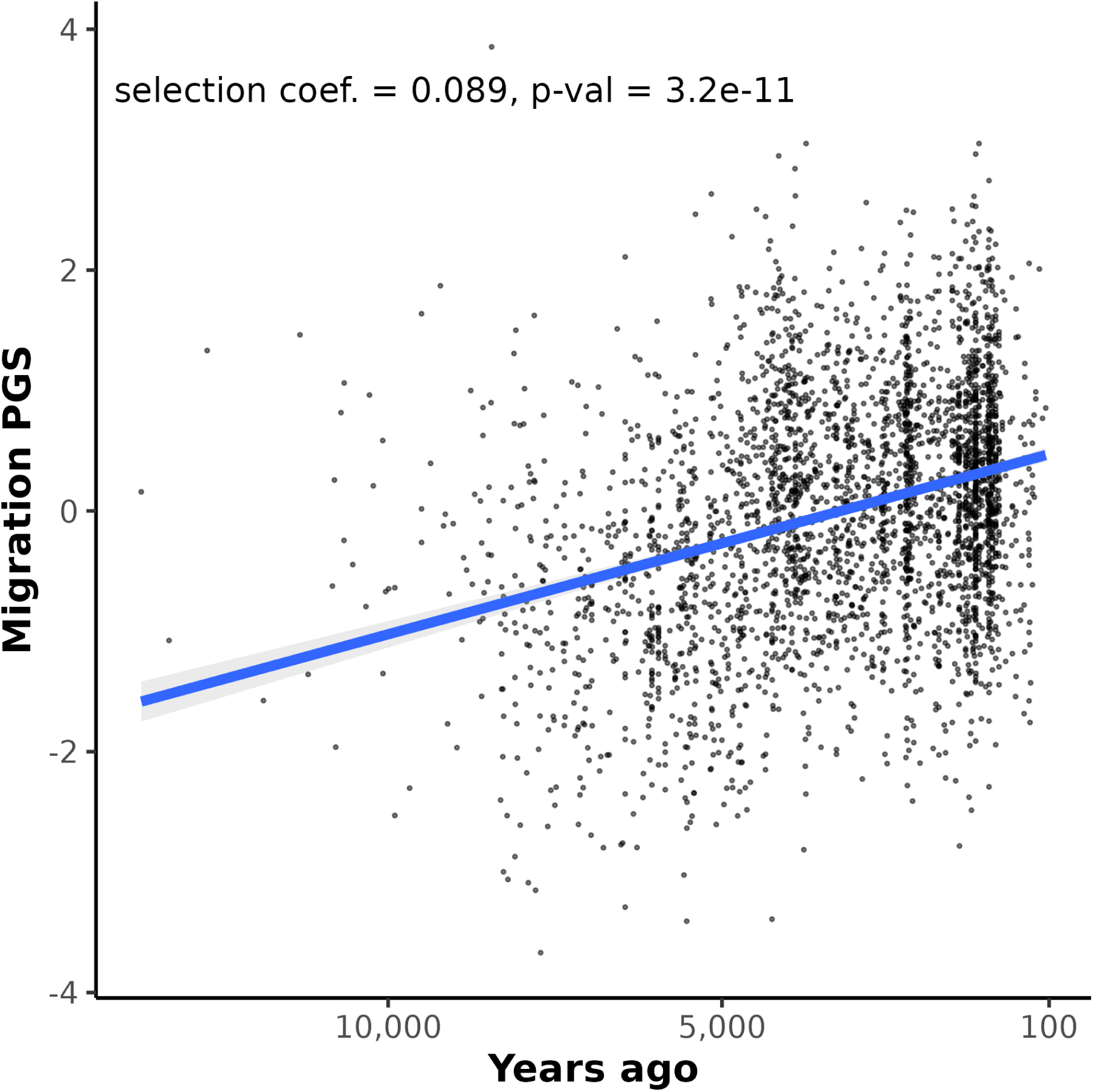
Selective pressures acting on migration alleles over the past 14,000 years. Analysis of the change in migration PGS over time from N = 3,241 ancient human genomes (X-axis = time before present, Y-axis = migration PGS). Statistics from a linear mixed model accounting for genetic relatedness and population structure are shown.

**Figure S3:**
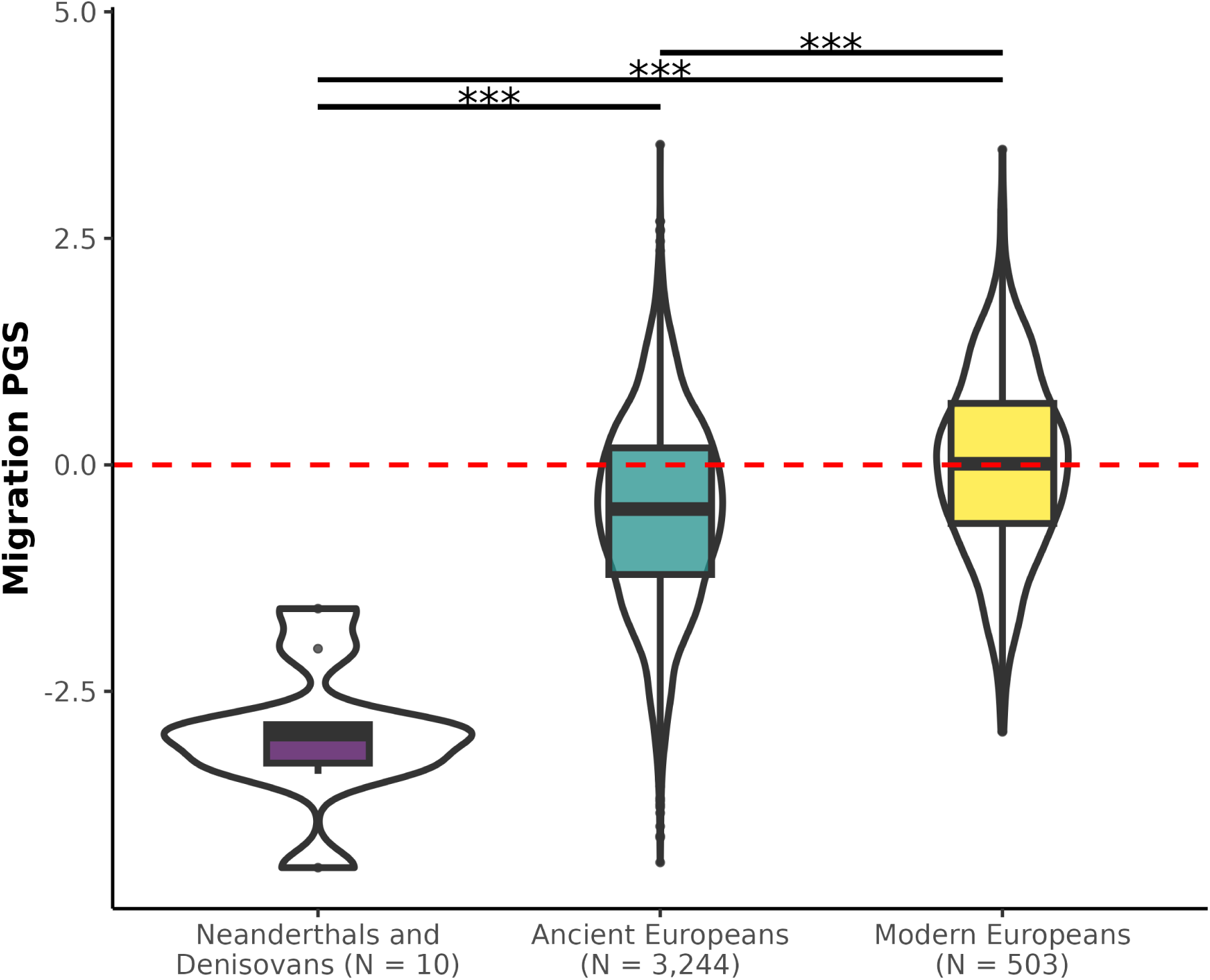
Comparison of migration PGS across hominins. *** indicates statistical significance (*p <* 0.001) from t-tests between groups.

**Figure S4:**
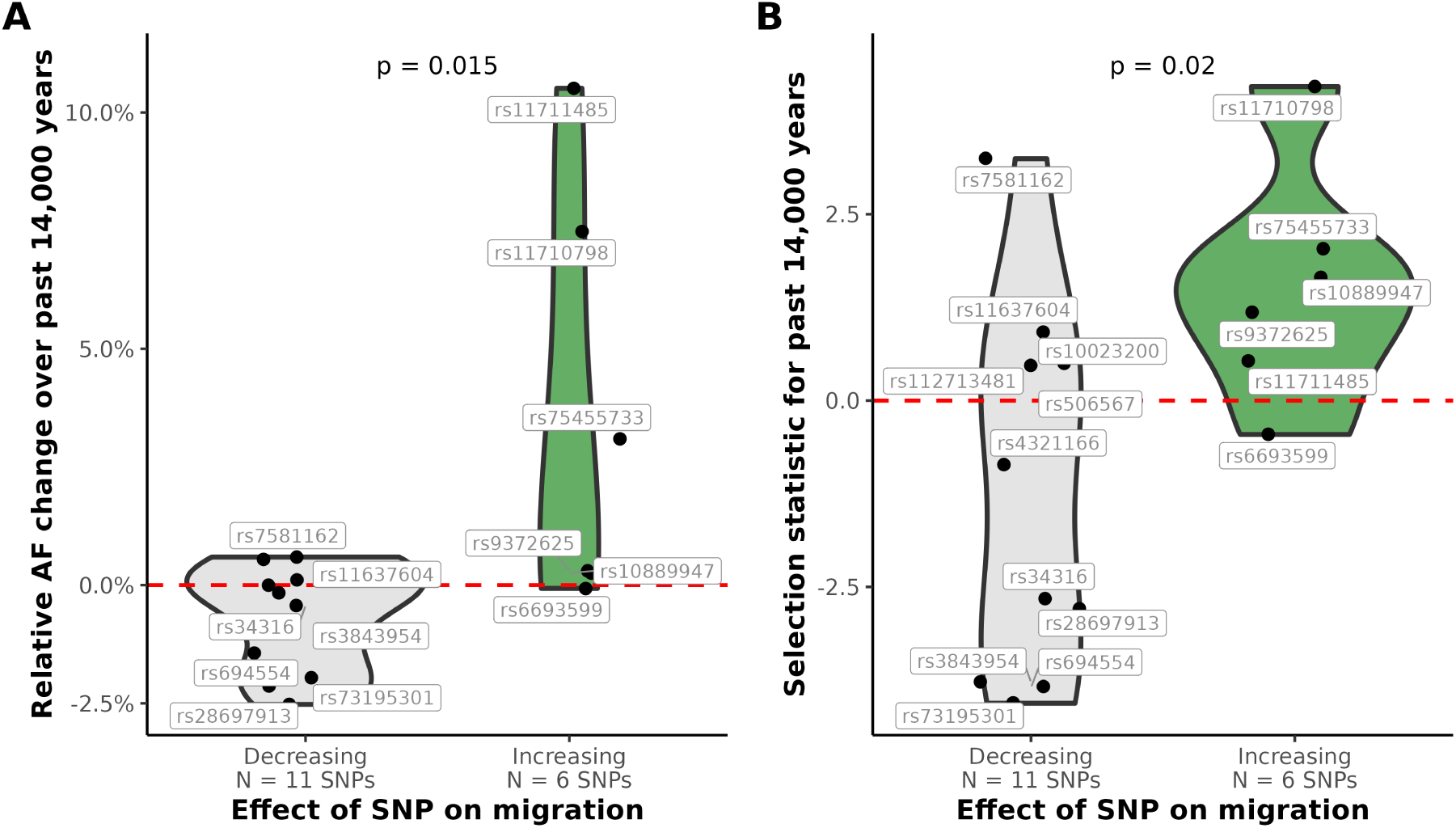
Evidence of directional selection at migration GWAS loci. **A** Comparing relative allele frequency changes over the past 14,000 years and whether that SNP is associated with increased or decreased migration in the UK Biobank. All SNPs analyzed are independent migration GWAS hits (*p <* 5 × 10^−8^). Allele frequency information came from the AADR (*36*). Relative AF change = allele frequency change / modern allele frequency. *p* from Wilcoxon test. **A** Comparing directional selection statistic over the past 14,000 years and whether that SNP is associated with increased or decreased migration in the UK Biobank. All SNPs analyzed are independent migration GWAS hits (*p <* 5 × 10^−8^). Selection statistics came from the AADR (*36*). *p* from Wilcoxon test.

**Figure S5:**
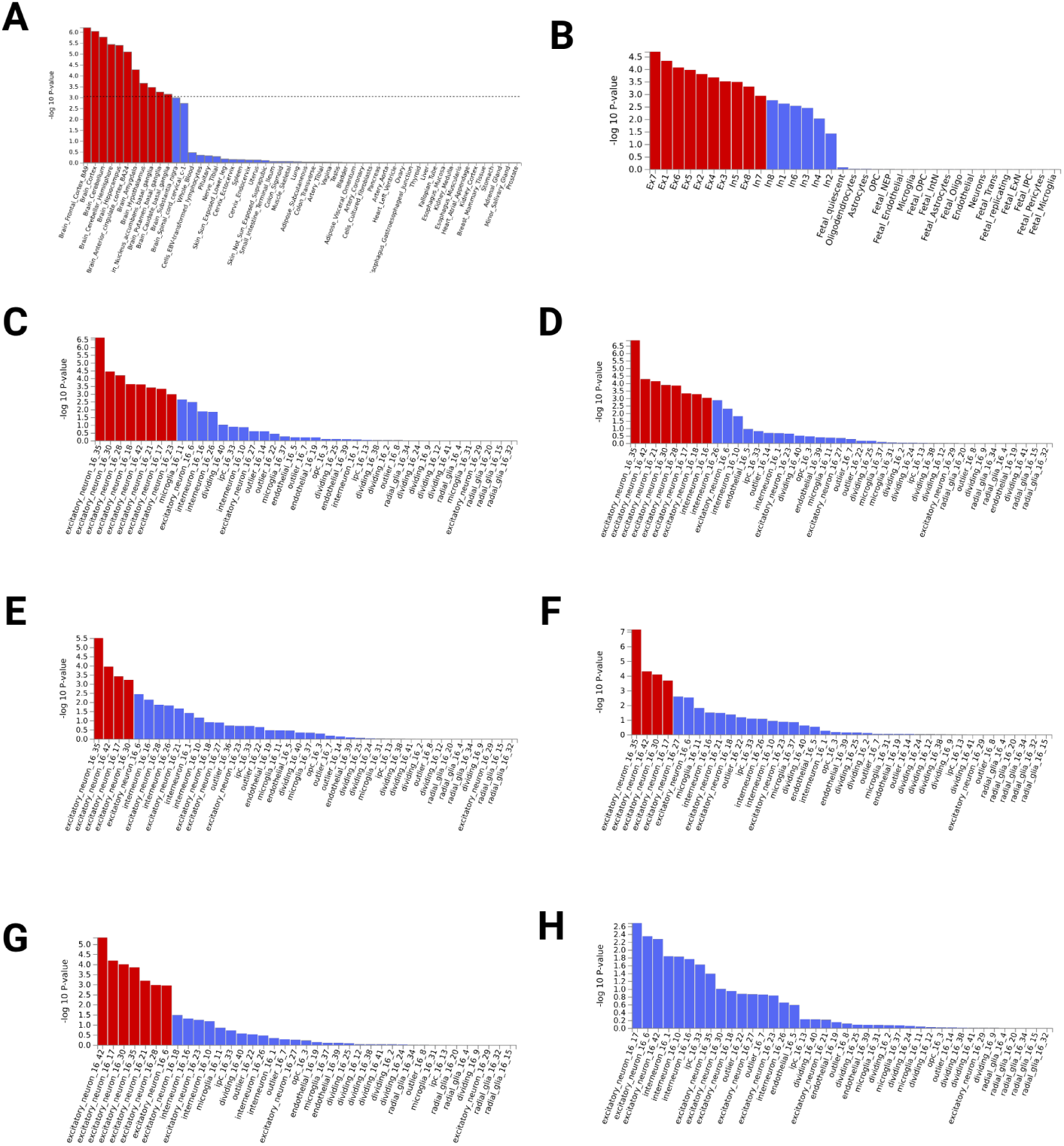
Enrichment of migration-associated genes across tissues and cell-types. Red bars indicates Bonferroni significant enrichment. **A** MAGMA enrichment of migration-associated genes across GTEx v8 tissues. **B** MAGMA enrichment of migration-associated genes across PsychENCODE developmental cell-types. **C** MAGMA enrichment of migration-associated genes across prenatal brain cell-types in the temporal cortex (16 weeks post conception). **D** MAGMA enrichment of migration-associated genes across prenatal brain cell-types in the parietal cortex (16 weeks post conception). **E** MAGMA enrichment of migration-associated genes across prenatal brain cell-types in the primary motor cortex (16 weeks post conception). **F** MAGMA enrichment of migration-associated genes across prenatal brain cell-types in the primary somatosensory cortex (16 weeks post conception). **G** MAGMA enrichment of migration-associated genes across prenatal brain cell-types in the primary visual cortex (16 weeks post conception). **H** MAGMA enrichment of migration-associated genes across prenatal brain cell-types in the prefrontal cortex (16 weeks post conception).

**Figure S6:**
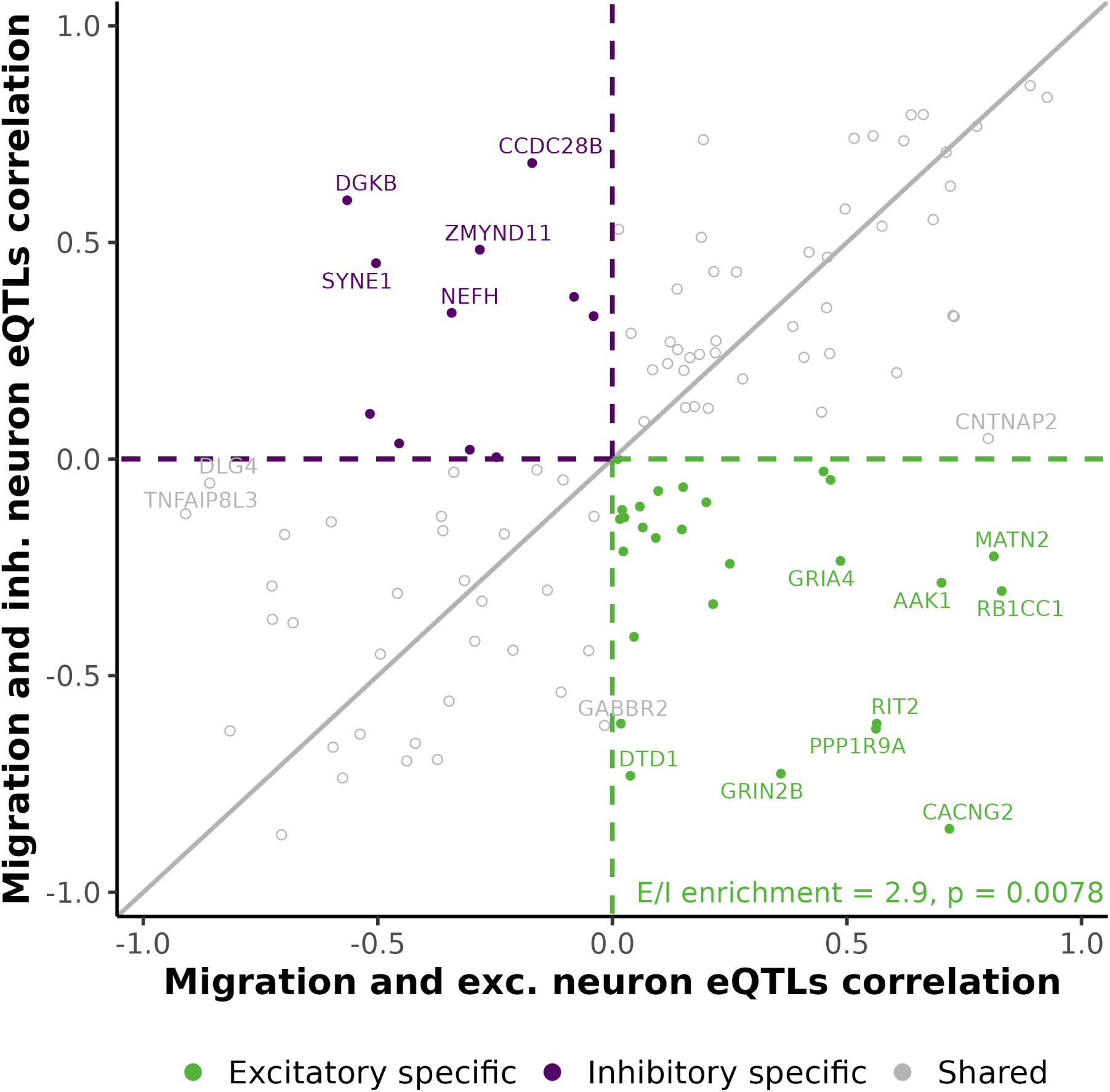
Migration-increasing alleles preferentially upregulate excitatory neuron markers. Comparison of excitatory versus inhibitory neuron specific effects of migration-associated variants. The axes represents gene level correlation coefficients between migration GWAS Z-scores and celltype specific eQTL Z-scores. The X-axis is the association between migration variant effects and excitatory neurons, and the Y-axis is the association with inhibitory neurons. Each point is a different neuronal marker gene identified in PanglaoDB.

**Figure S7:**
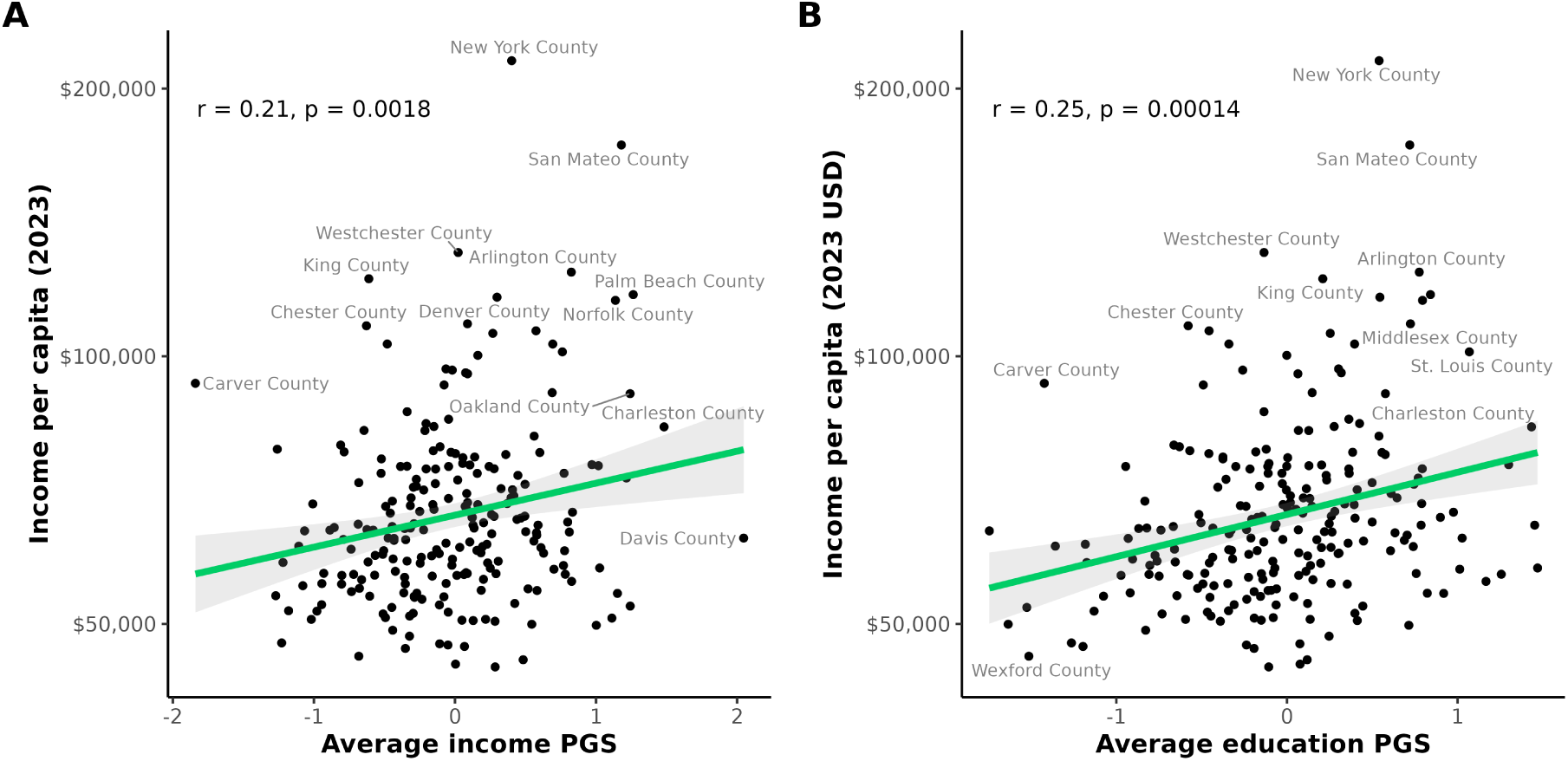
US county-level associations between average polygenic scores and current income. (**A**) Relationship between county-level average income polygenic scores (PGS) and per-capita income in 2023. (**B**) Relationship between county-level average educational attainment PGS and per-capita income in 2023. Each point represents one county, with the solid line indicating the linear regression fit and shaded area showing the 95% confidence interval. Pearson’s correlation coefficient (*r*) and corresponding *p* are reported in each panel.

**Figure S8:**
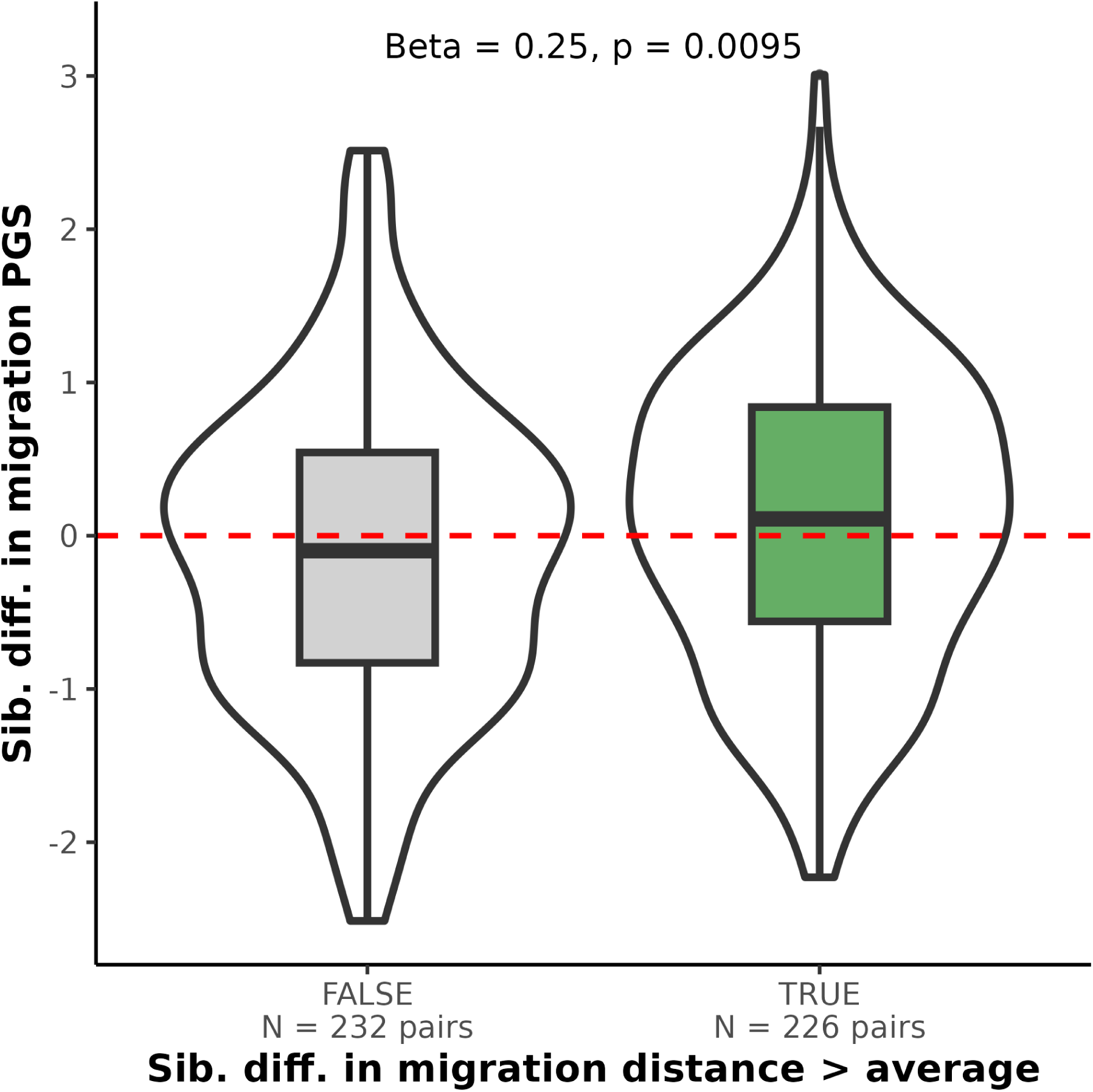
Siblings with larger differences in migration distance have more migration PGS differences. Sibling pairs were split into two groups, based on whether or not they migrated further from each other than the median distance. Logistic regression results are shown, regression model accounted for differences in age, sex, birth year, genetic PCs 1-5, and education level.

## Notes

https://www.ukbiobank.ac.uk

https://base.sfari.org/

https://www.internationalgenome.org/data/

https://reich.hms.harvard.edu/allen-ancient-dna-resource-aadr-downloadable-genotypes-present-day-and-ancient-dna-data

### Competing Interest Statement

The authors have declared no competing interest.

